# *Stenotrophomonas maltophilia* promotes wheat growth by enhancing nutrient assimilation and rhizosphere microbiota modulation

**DOI:** 10.1101/2025.02.26.640485

**Authors:** Pinki Sharma, Rajesh Pandey, Nar Singh Chauhan

**Author notes:** Corresponding author **Nar Singh Chauhan**, **Rajesh Pandey**.

## Abstract

**Background:** *Stenotrophomonas maltophilia* has gained considerable attention for its biocontrol and biofertilizer potential in promoting plant growth. It could be employed to enhance wheat yield to ensure food security for the growing population. However, its biofertilizer potential in field conditions and its impact on wheat rhizosphere microbiota must be assessed before its employment in agriculture practices to increase wheat production.

**Methods:** We have assessed the role of *Stenotrophomonas maltophilia* on wheat seed germination, plant growth parameters, and crop yield in the field conditions. Additionally, wheat rhizosphere microbiota was explored to assess the impact of seed pretreatment with *Stenotrophomonas maltophilia* on the wheat rhizosphere microbiota.

**Results and Discussion:** *Stenotrophomonas maltophilia* strains BCM and BCM_F demonstrated superior antifungal activity, indicating their biocontrol potential. Seed pretreatment with these strains promoted nitrogen fixation and phosphate solubilization in the wheat rhizosphere showcasing biofertilizer potential. Uniquely identified OTUs in the rhizosphere microbiota of treated groups and microbial community dynamics, particularly at Feeks 3.0 and 6, indicated *Stenotrophomonas maltophilia*- induced microbiota restructuring. The abundance of *Stenotrophomonas maltophilia* 16S rRNA gene sequences at different Feeks treated with microbial indicates its stability across different plant growth stages. Their rhizospheric presence also impacted plant health indicators, including improved sugar and nitrite concentrations and significantly enhanced crop yield (*P>0.05*). Enhanced growth parameters and better crop yield in *Stenotrophomonas maltophilia* pre-inoculated seeds in field conditions indicated their potential to offer a sustainable alternative to enhance wheat production.

**Conclusion:** The present study highlighted the biofertilizer and biocontrol potential of *Stenotrophomonas maltophilia* strains BCM and BCM_F in supporting sustainable agricultural practices.

## Introduction

*Stenotrophomonas maltophilia* is a gram-negative, motile, non-fermentative bacterium identified from various habitats like soil, water, plants, and animal hosts. It has gained considerable attention for its plant growth potential. *S. maltophilia* produces siderophores (*ent*A, *ent*B, *ent*C, *ent*D, *ent*E) and spermidine (*spe*A, *spe*C) extending the beneficial effects on plant growth and development (Ulrich et al., 2021). *Stenotrophomonas* sp. could fix atmospheric nitrogen in *Oryza sativa* and *Glycine max* via *nif*H and related gens expression (Alexander et al., 2019). *S. maltophilia* harbors various phosphate solubilization genes (*pstC, pstA, pstS, phoU*), allowing phosphate solubilization from inorganic phosphate minerals. These nutrient-assimilation abilities are particularly beneficial for crops grown in nutrient-poor soils. Furthermore, *S. maltophilia* produces a variety of phytohormones, including auxins, cytokinins, and gibberellins, which are known to promote root development, seed germination, and overall plant growth (Kumar et al., 2023). The production of these phytohormones enhances root architecture, facilitating better nutrient uptake. In wheat (*Triticum aestivum*), inoculation with *Stenotrophomonas maltophilia* has resulted in increased root biomass and improved nutrient absorption, demonstrating its role in enhancing root vigor and facilitating better growth in nutrient-limited conditions (Khoso et al., 2024). *Stenotrophomonas maltophilia* also contributes to plant health by exhibiting antagonistic activity against phytopathogens. *Stenotrophomonas maltophilia* suppressed the Fusarium wilt in *Solanum lycopersicum* and *Solanum bonariense,* caused by *Fusarium oxysporum* (Abdallah et al., 2020). *Stenotrophomonas maltophilia* has also shown a protective effect against *Ralstonia solanacearum,* the causative agent of potato brown rot*. Stenotrophomonas maltophilia* UPMKH2 foliar spray inhibits the rice blast disease caused by the *Pyricularia oryzae* (Messiha et al., 2007)*. S. maltophilia* SBP-9 has been reported to develop biotic and abiotic resistance in various crops (Singh and Jha, 2017). *Stenotrophomonas maltophilia* BCM was also characterized for biofertilizer and biocontrol potential in wheat against two phytopathogens, *Rhizoctonia solani* and *Fusarium oxysporum* (Sharma et al., 2024a). However, these properties still need to be assessed in field conditions. Additionally, the stability of *Stenotrophomonas maltophilia* in the wheat rhizosphere after seed pretreatment and the impact of pre-inoculation on wheat rhizosphere microbiota still awaits exploration. The current study analyzed the plant growth promotion potential of *S. maltophilia* under field conditions in addition to assessing the impact of *S. maltophilia* pre-inoculation on wheat rhizosphere microbiota at different plant growth stages.

## Methodology

### Site description and sampling

Rhizospheric samples were obtained from wheat plants grown in an experimental field located in the botanical garden of Maharshi Dayanand University, Rohtak, Haryana, India (28° 52’ 44’’ N, 76° 37’ 19’’ E). Wheat rhizospheric microbes were isolated using previously standardized methods (Sharma et al., 2024a).

### Molecular, physiological, and biochemical characterization of microbial isolate BCM_F

Gram staining of the biocontrol microbe was performed using a Gram staining kit (Himedia, K001-1KT). The growth of the microbe was observed at various pH (3.0 to 12.0) and temperature (10 to 60°C) ranges to determine optimal conditions. Growth patterns in LB broth were monitored for 48 hours at 37°C with shaking at 200 rpm to assess its doubling time (Wang et al., 2015). Substrate utilization was evaluated using the Hi Carbo kit (Himedia). Biochemical properties were analyzed through amylase, catalase, pectinase, cellulase, esterase, and protease assays. Antibiotic susceptibility was tested using Combi IV and G-VI-plus kits (Himedia). Stress response was evaluated by testing for salt, metal, and oxidative stress tolerance (Sharma et al., 2024). Drought stress tolerance of *Stenotrophomonas maltophilia* BCM and *Stenotrophomonas maltophilia* BCM_F was assessed in the presence of PEG as a drought stressor (Sharma et al., 2024a, b). ACC deaminase production activity of microbial isolates was checked with a standardized methodology (Maheshwari et al., 2020). ACC deaminase activity was calculated based on the amount of α-Ketoglutarate in the reaction mixture using the α- Ketoglutarate standard curve (R=0.9998). DNA extraction was done via the alkali lysis method (Chauhan et al., 2009), with qualitative and quantitative analysis using agarose gel electrophoresis and Qubit HS DNA kits (Invitrogen). The 16S rRNA gene was amplified and sequenced for taxonomic identification (Sharma et al., 2024a).

### Role of *Stenotrophomonas maltophilia* BCM and BCM_F on seed germination under phytopathogen and saline stress condition

The biocontrol potential of *Stenotrophomonas maltophilia* BCM and *maltophilia* BCM_F was evaluated against two plant pathogenic fungi, *Rhizoctonia solani* and *Fusarium oxysporum*. The assessment focused on their impact on seed germination efficiency and root and shoot length of wheat seedlings (Sharma et al., 2024b, Singh and Kayastha, 2014). To assess the role of *Stenotrophomonas maltophilia* BCM and BCM_F on seed germination under saline conditions, seeds were initially soaked in an overnight-grown microbial culture containing 10^11^cells/ml, with varying concentrations of NaCl ranging from 0 to 1M, for 16 hours at 37°C.

Control seeds were soaked directly in NaCl solutions of the same concentrations for 16 hours at 37°C. After soaking, the seeds were wrapped in germination sheets and placed in 50 ml culture tubes containing 5 ml of Hoagland solution. They were then incubated in the dark at ambient temperature (25° C) for 7 days. Wheat seed germination percentage, alpha-amylase activity, and root and shoot lengths were measured after the incubation (Sharma et al., 2024b, Singh and Kayastha, 2014).

### Biofertilizer potential of *Stenotrophomonas maltophilia* in experimental field conditions

Roots of WC-306 plants treated with *Stenotrophomonas maltophilia* BCM and BCM_F were harvested at different Feeks and analyzed for phosphate-solubilizing ability (Behera et al., 2017), nitrate reductase activity (Kim and Seo, 2018), total sugar content (Ludwig and Goldberg, 1956), and reducing sugar content (Khatri and Chhetri, 2020), compared to untreated WC-306 plant. In addition, various growth and yield parameters were evaluated in both treated and untreated plants, including the number of tillers per plant, number of leaves per plant, number of spikes per plant, spike length, number of spikelets per spike, and grain yield, to assess the impact of *Stenotrophomonas maltophilia* BCM and BCM_F on plant growth and productivity.

### Wheat rhizosphere microbiota profiling

Seeds of the wheat cultivar 306, treated with microbial isolates, were cultivated in an experimental field at the botanical garden of Maharshi Dayanand University, Rohtak, Haryana, India (28° 52’ 44’’ N, 76° 37’ 19’’ E). Wheat roots were harvested at different Feeks (1.0, 2.0, 3.0, 6.0, 9.0, and 10.5) (https://www.sunflower.k-state.edu/agronomy/wheat/wheatdevelopment.html). Metagenomic DNA from the wheat rhizosphere was extracted using the CTAB method as described by Kumar et al. 2016 (Kumar et al., 2016), followed by purification with the HiPurA soil DNA purification kit (Himedia). The 16S rRNA gene sequences were amplified from the wheat rhizosphere metagenomic DNA using universal primers (27F 5’-AGAGTTTGATCCTGGCTCAG-3’; 1492R 5’- GGTTACCTTGTTACGACTT-3’) (Sharma et al., 2024b). The 16S rRNA gene amplicons were sequenced using the Nanopore MinION MK1C with the Midnight pipeline. The resulting wheat rhizosphere data were analyzed using EPIME 16S rRNA gene pipelines following their default settings (Sharma et al., 2024b).

### Statistical analysis

All experiments were conducted in replicates. Statistical analyses and graphical representations of the datasets were performed using SIGMA Plot 15. ANOVA was used to determine the significance between microbial-treated and non-treated groups [Systat Software SigmaPlot 15].

## Results

### Isolation and screening of microbes with biocontrol potential

The wheat rhizospheric samples have a pH of 7.3±0.0025, temperature 22.6°C±0.0102 and moisture content (11.5±1.10014). Twelve morphologically diverse microbes were purified from the wheat rhizosphere, however, only two isolates (BCM and BCM_F) showcased antifungal activity. The microbial isolate BCM produced growth inhibition zones of 17±0.57 mm against *Rhizoctonia solani* and 15±0.57 mm against *Fusarium oxysporum*. In contrast, the microbial isolate BCM_F demonstrated a more potent antifungal effect, achieving a growth inhibition zone of 22±0.2 mm specifically against *Fusarium oxysporum*. These findings highlight the potential of both microbial isolates in suppressing fungal growth, with BCM_F showing particularly significant efficacy against *Fusarium oxysporum*, which may suggest their potential in biological control strategies for managing fungal infestation, especially during seed germination.

### Molecular, physiological, and biochemical characterization of microbial isolates

The 16S rRNA gene of the microbial isolate BCM_F showed 99.38% homology with *Stenotrophomonas maltophilia* IAM 12423 (NR 041577.1). Phylogenetic analysis further supported its affiliation with *Stenotrophomonas maltophilia* (**Figure 1A**). Microscopic examination revealed that BCM_F is a gram-negative, rod-shaped, motile bacterium. It exhibited optimal growth at pH 7.0 and 37°C and reached the log phase after 16 hours with a doubling time of approximately 42.86 minutes (**Figure 1B**). BCM_F displayed facultative anaerobic growth, with an O.D. of 0.506 at 600nm after 24 hours under anaerobic conditions. Biochemically, BCM_F was positive for amylase, esterase, lipase, protease, and catalase activities. Its substrate utilization profile was similar to other *Stenotrophomonas maltophilia* species. Antibiotic susceptibility testing showed resistance to several antibiotics, including bacitracin, cephalothin, vancomycin, ofloxacin, and erythromycin, while being sensitive to others like amikacin, ceftazidime, and lincomycin. The antibiotic resistance profile resembled other *Stenotrophomonas maltophilia* strains, especially *Stenotrophomonas maltophilia* smyn 45. Stress response assays demonstrated BCM_F’s ability to grow in the presence of various salts (up to 4.93% NaCl (w/v), up to 9.2% KCl (w/v), up to 4.4% LiCl (w/v)), metals (up to 0.1% Na_3_AsO_4_ (w/v),0.1% NaAsO_2_ (w/v), and 0.3% CdCl_2_ (w/v)), and oxidizing agents (up to 6.11% (v/v) H_2_O_2_). The microbial isolates BCM and BCM_F exhibited robust growth up to 40% of polyethylene glycol (PEG) concentration (**Figure 1C**). Interestingly, BCM_F showed greater activity than BCM. These findings highlight the plant growth-promoting features under drought stress. Microbial isolate BCM_F (0.332 EU) was found to produce more ACC deaminase enzyme than BCM (0.299 EU) (**Figure 1D**). ACC deaminase activity in both strains confirms their oxidative stress-mitigating properties. The molecular, physiological, and biochemical characterization of microbial isolate BCM has identified it as a gram-negative, rod-shaped, motile, facultative aerobic bacterium having a generation time of 67.8 min in optimum growth conditions (pH 7.0 and Temperature 35°C) (Sharma et al., 2024a). The phylogenetic and phylogenomic analysis confirmed it as the strain of *Stenotrophomonas maltophilia* (Sharma et al., 2024a).

**Figure 1:**
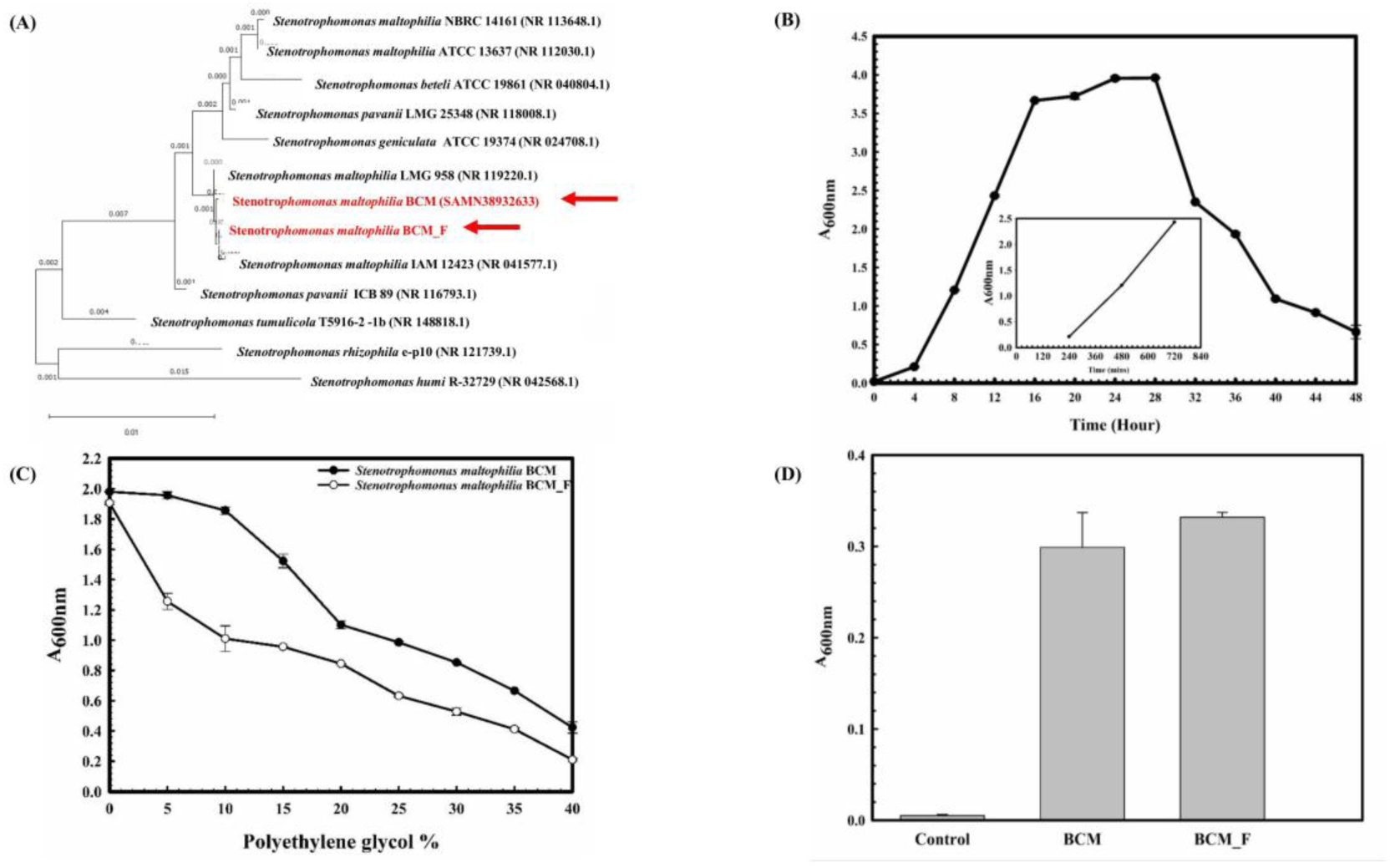
Phylogenetic, Physiological, and biochemical characterization of *Stenotrophomonas maltophilia* BCM and BCM_F. Phylogenetic affiliation of microbial isolates *Stenotrophomonas maltophilia* BCM and BCM_F with the other *Stenotreophomonas* species **(A)**. The phylogenetic tree was constructed using the Neighbor-joining method of phylogenetics with 1000 bootstrap replications using MEGA-X software. The out-group was represented by *Stenotrophomonas humi* R-32729 SSU rRNA gene sequence. Growth pattern analysis of *Stenotrophomonas maltophilia* BCM_F was observed after incubating the cultures for 48 hours in LB broth with constant shaking at 200 rpm **(B)**. The experiment was carried out in triplicates, and growth was observed by taking absorbance at 600 nm after 4 hours. The drought stress tolerance ability of *Stenotrophomonas maltophilia* BCM and BCM_F **(C)**. Bacterial growth was observed after incubating the cultures in nutrient broth supplemented with different PEG concentrations at 37°C for 24 h with constant shaking at 200 rpm. The experiment was carried out in triplicates, and growth was observed by taking absorbance at 600 nm. ACC deaminase activity of *Stenotrophomonas maltophilia* BCM and BCM_F (D). Microbial cultures were incubated with ACC, and the resulting ammonia concentration was quantified using a colorimetric method. Values plotted are the triplicate readings’ mean and the observed standard deviation.

### *Stenotrophomonas maltophilia* BCM and BCM_F role on seed germination under phytopathogen exposure and saline stress condition

The control group exhibited a germination percentage of 75.66±0.57735. However, seeds pre-treated with *Stenotrophomonas maltophilia* BCM and BCM_F showed germination rates of 96.67±0.5% and 93.33±0.57%, respectively (**Figure 2A**). This corresponds to >100 fold increased seed germination. In the presence of *Rhizoctonia solani* and *Fusarium oxysporum*, seed germination was reduced to 10±1% and 5±0.57% respectively. Pre-treatment with *Stenotrophomonas maltophilia* BCM significantly increased the germination percentage to 48.52±0.57 and 45.66±0.57 in the presence of *Rhizoctonia solani* and *Fusarium oxysporum*, respectively (**Figure 2A**). It was significantly higher than the untreated control groups incubated with *Rhizoctonia solani* (*P=0.0015*) and *Fusarium oxysporum* (*P=0.024*). Similarly, seeds pre-treated with *Stenotrophomonas maltophilia* BCM_F showed increased seed germination to 13.56±0.02% and 53.52±0.70% in the presence of *Rhizoctonia solani* and *Fusarium oxysporum*, respectively. It was significantly higher than the untreated control groups exposed to *Rhizoctonia solani* (*P=0.045*) and *Fusarium oxysporum* (*P=0.003*). *Stenotrophomonas maltophilia* BCM and BCM_F significantly increased seed germination in the presence of *Fusarium oxysporum* by ∼ 913-fold and 1070-fold, respectively, compared to untreated control seeds. Similarly, higher seed germination (∼480-fold) was observed for *Stenotrophomonas maltophilia* BCM pre-inoculated seeds in the presence of *Rhizoctonia solani* compared to the un-inoculated control. These results strongly indicate the biocontrol potential of both microbial strains. *Rhizoctonia solani* and *Fusarium oxysporum* exposure to wheat seeds significantly reduced alpha-amylase activity (*P=0.0001* and *P=0.0018*) in wheat seeds, respectively. Pre-treatment with *Stenotrophomonas maltophilia* BCM and BCM_F significantly increased alpha-amylase activity in the presence of *Rhizoctonia solani* (*P=0.00013* and *P=0.00001* respectively) and *Fusarium oxysporum* (*P=0.0001* and *P=0.00003* respectively) (**Figure 2B**). The significant increase in alpha-amylase activity following pre-treatment with *Stenotrophomonas maltophilia* BCM and BCM_F could explain their role in enhanced seed germination.

**Figure 2:**
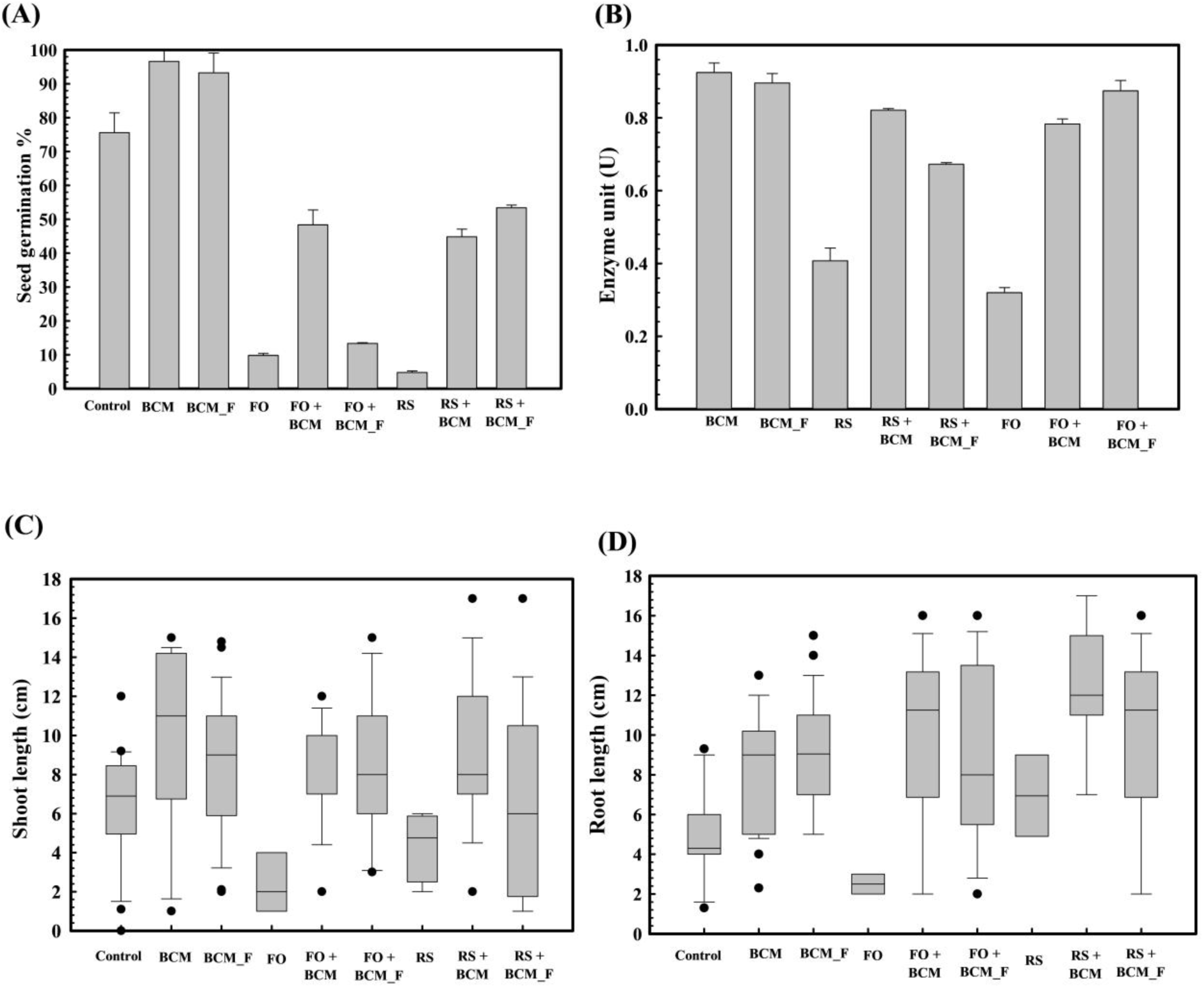
Effect of *Stenotrophomonas maltophilia* BCM and BCM _F on wheat seed germination parameters. The seed germination percentage in *Stenotrophomonas maltophilia* BCM and BCM _F pretreated seed in reference to the control (untreated) (**A**). The alpha-amylase activity of *Stenotrophomonas maltophilia* BCM and BCM _F pretreated wheat seeds in the presence and absence of plant pathogenic fungi *Rhizoctonia solani* and *Fusarium oxysporum* (**B).** The impact of seeds pretreatment with *Stenotrophomonas maltophilia* on shoot **(C)** and root length **(D)** of WC-306 seedling in the presence of *Rhizoctonia solani* and *Fusarium oxysporum* (**C).** Plotted values are the mean of triplicates along with the observed standard deviation. Here BCM: *Stenotrophomonas maltophilia* BCM; BCM_F: *Stenotrophomonas maltophilia* BMC_F; C: control; FO: *Fusarium oxysporum*; RS: *Rhizoctonia solani*.

Pre-treatment of wheat seeds with *Stenotrophomonas maltophilia* BCM and BCM_F not only enhanced seed germination but also improved the growth of wheat plantlets. Seeds treated with *Stenotrophomonas maltophilia* BCM and BCM_F exhibited significantly increased shoot length (*P=0.027*, *P=0.0023*) and root length (*P=0.010*, *P=0.024*) compared to the untreated seeds (**Figure 2C**). Additionally, *Stenotrophomonas maltophilia* BCM significantly improved the root (*P=0.020*, *P=0.032*) and shoot length (*P=0.031*, *P=0.129*) of plantlets obtained from the wheat seeds exposed with *Rhizoctonia solani* and *Fusarium oxysporum*, respectively. Similarly, *Stenotrophomonas maltophilia* BCM_F significantly enhanced the root (*P=0.0027*, *P=*0.019) and shoot length (*P=*0.020, *P=*0.039) of plantlets arising from wheat seeds exposed with *Fusarium oxysporum* (**Figure 2D**). Despite the exposure to *Rhizoctonia solani* and *Fusarium oxysporum,* the average shoot and root lengths of wheat plantlets treated with Stenotro*phomonas maltophilia* BCM and BCM_F, remained significantly higher (*P>0.05*) compared to the control, highlighting their biocontrol potential.

The role of *Stenotrophomonas maltophilia* BCM and BCM_F in wheat seed germination under salinity stress was also evaluated. Seed germination was significantly reduced with increased salt concentration (**Figure 3**). However, seeds pre-treated with *Stenotrophomonas maltophilia* BCM and BCM_F exhibited improved germination under high salinity conditions (**Figure 3)**. An approximately 12.85- and 7.78-fold increased seed germination was observed with seeds pre-treated for BCM and BCM_F, respectively, even at the higher saline conditions like 1M NaCl concentration, compared to control. Pre-treatment with these microbial strains not only enhanced seed germination but also promoted the growth of wheat plantlets (**Figure 4**). Seeds treated with *Stenotrophomonas maltophilia* BCM and BCM_F showed significantly increased shoot length (**Figure 4C, E**) (*P=0.001* and *P=0.0001*) and root length (**Figure 4D, F**) (*P=0.001* and *P=0.0003*) compared to seedling germinated from the untreated seeds (**Figure 4 A, B**) under high salinity conditions, respectively.

**Figure 3:**
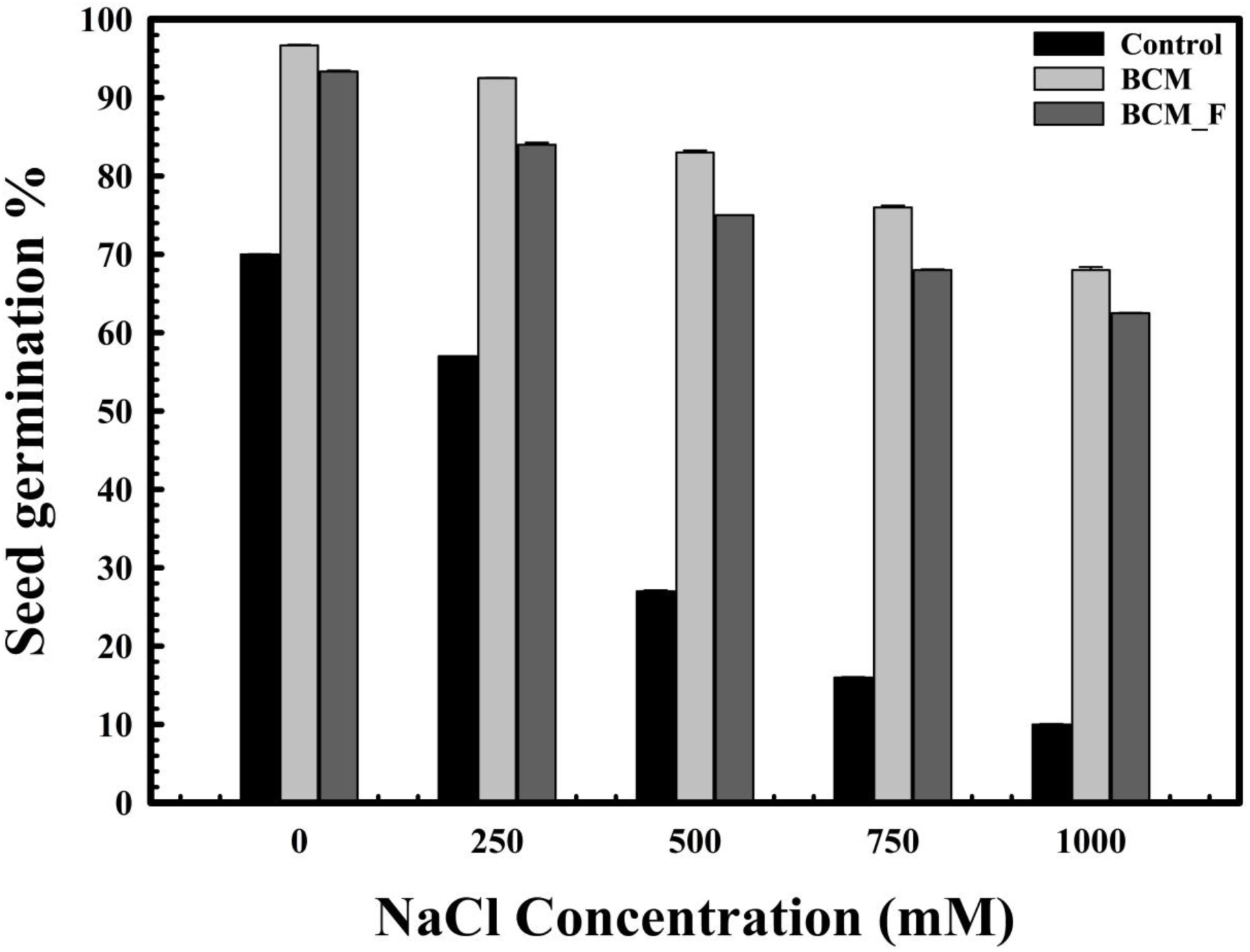
The impact of *Stenotrophomonas maltophilia* pre-treatment on wheat seed germination in saline conditions. The seed germination assays were performed to compare the effect of *Stenotrophomonas maltophilia* BCM and BCM _F isolates on seed germination under saline conditions. All assays were performed in triplicates.

**Figure 4:**
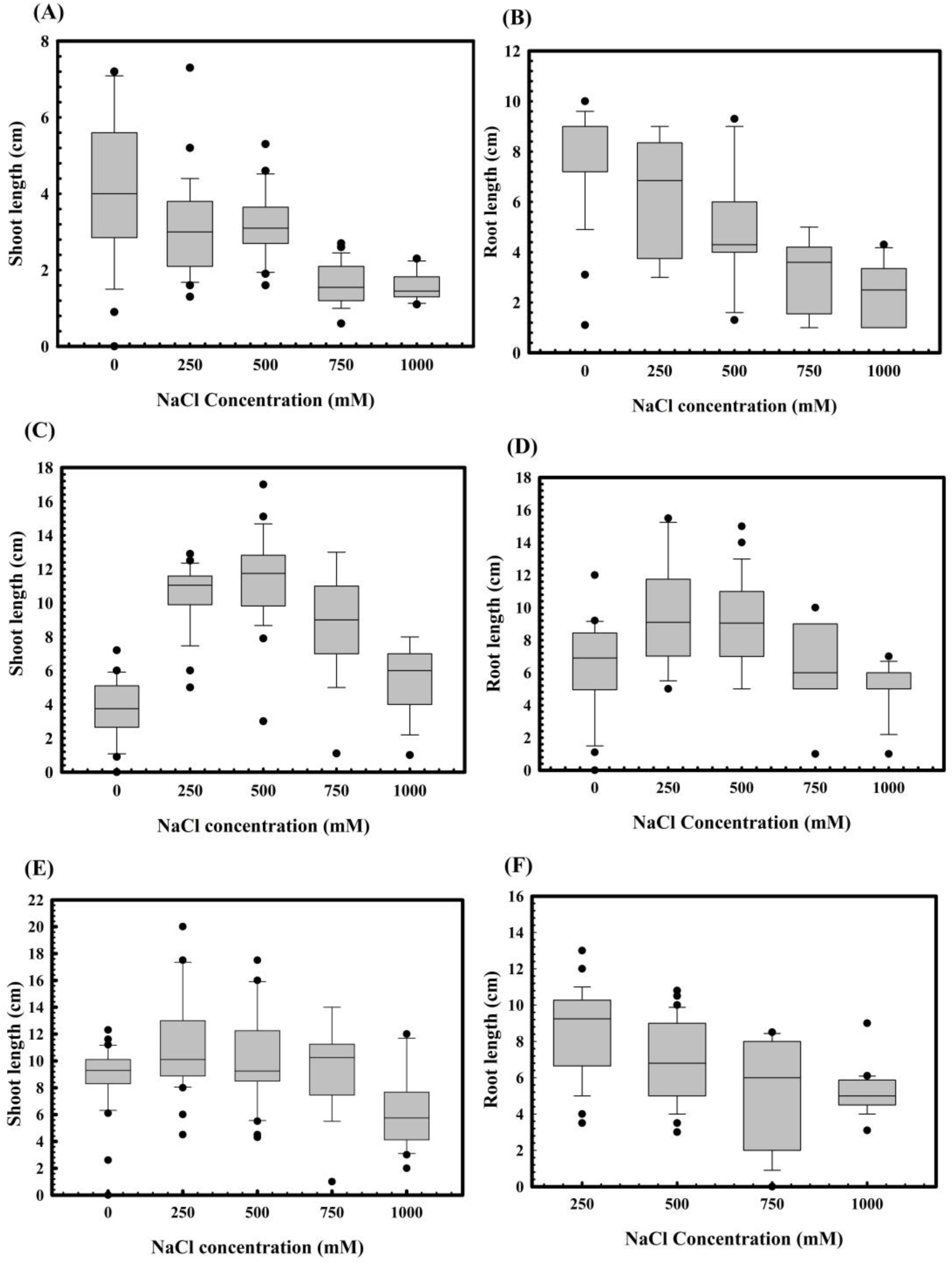
Effect of salinity stress on the root and shoot lengths of WC-306 plantlets. Shoot **(A)** and root **(B)** length of WC-306 seedlings under NaCl induced salinity stress. Shoot **(C)** and root **(D)** lengths of WC-306 seedlings of *Stenotrophomonas maltophilia* BCM pre-treatment group under NaCl-induced salinity stress. Shoot **(E)** and root **(F)** lengths of WC-306 seedlings of *Stenotrophomonas maltophilia* BCM_F pre-treatment group under NaCl-induced salinity stress. All assays were performed in triplicates.

### Wheat rhizosphere microbiota profiling

Rhizosphere microbiota was assessed at six Feeks to assess microbial community dynamics across plant growth stages. These Feeks were selected based on their significance in plant growth and development. Feeks 1 (emergence) is crucial for assessing whether microbial inoculants improve seed germination and early seedling growth, especially under challenging soil conditions or stress factors. Feeks 2, the beginning of tillering, is essential in studying the effects of inoculation on vegetative growth. Microbes that promote nutrient availability or enhance plant growth may initiate more tillers, potentially leading to greater biomass and grain yield. The tillering formation (Feeks 3) allowed us to evaluate that how microbial inoculants influence the development of lateral shoots, which contribute directly to yield potential by increasing the number of productive stems. Feeks 6 corresponds to internode formation and is significant for understanding the influence of microbial treatments on stem elongation and overall plant structure. Microbes that improve nutrient uptake or facilitate more efficient use of resources can lead to taller, healthier plants with better light interception, contributing to improved photosynthesis and growth. Feeks 9, when the ligule of the leaves becomes visible, marks the transition from vegetative to reproductive growth. These Feeks are critical for studying microbial impacts on plant preparations for reproduction, including enhancing nutrient uptake during the crucial phases leading to flowering and grain filling. Finally, Feeks 10.5, heading complete, is particularly important for knowing the reproductive success of crops. Heading refers to the emergence of the flower head, and at this stage, the plant is preparing for pollination and grain formation. Microbial inoculations may influence flowering synchronization, pollination success, or post-heading stress tolerance, which are significant factors in determining final grain yield and quality. These stages serve as benchmarks for assessing the effectiveness of microbial inoculants in enhancing growth, yield, and stress tolerance. Metagenomic analysis of rhizosphere microbiota at different wheat growth stages revealed variations in the microbiota with wheat growth stages. Proteobacteria was the most abundant phylum across all studied Feeks, regardless of whether the samples were treated or from the non-treated control group. However, its abundance was variable across all Feeks. Notably, Feeks 3.0 and Feeks 6.0 exhibited a higher abundance of Proteobacteria than the other Feeks (**Supplementary Figure SF1**). Furthermore, the abundance of various proteobacterial classes also varied with Feeks (**Supplementary Figure SF1**). Microbial diversity analysis at the genus level has enriched our understanding of microbial community dynamics.

### Wheat rhizosphere microbiota composition at Feeks 1.0

At Feeks 1.0 (emergence stage), 449 operational taxonomic units (OTUs) were identified. The shared OTUs accounted for 5.79% of total diversity. Shared OTUs between control and BCM comprised 20.04% of total diversity, while those between BCM and BCM_F comprised 13.5%. Shared OTUs between BCM_F and control represented 2.44%. Unique OTUs in BCM, BCM_F, and the untreated control represented 20.93%, 23.6%, and 13.58% of total diversity, respectively (**Figure 5A**). Taxonomic affiliation of these OTUs identified shared and unique microbial taxa groups associated with various test groups (**Table 1**). These distinctions help to understand the microbial diversity and dynamics under the different experimental conditions.

**Figure 5:**
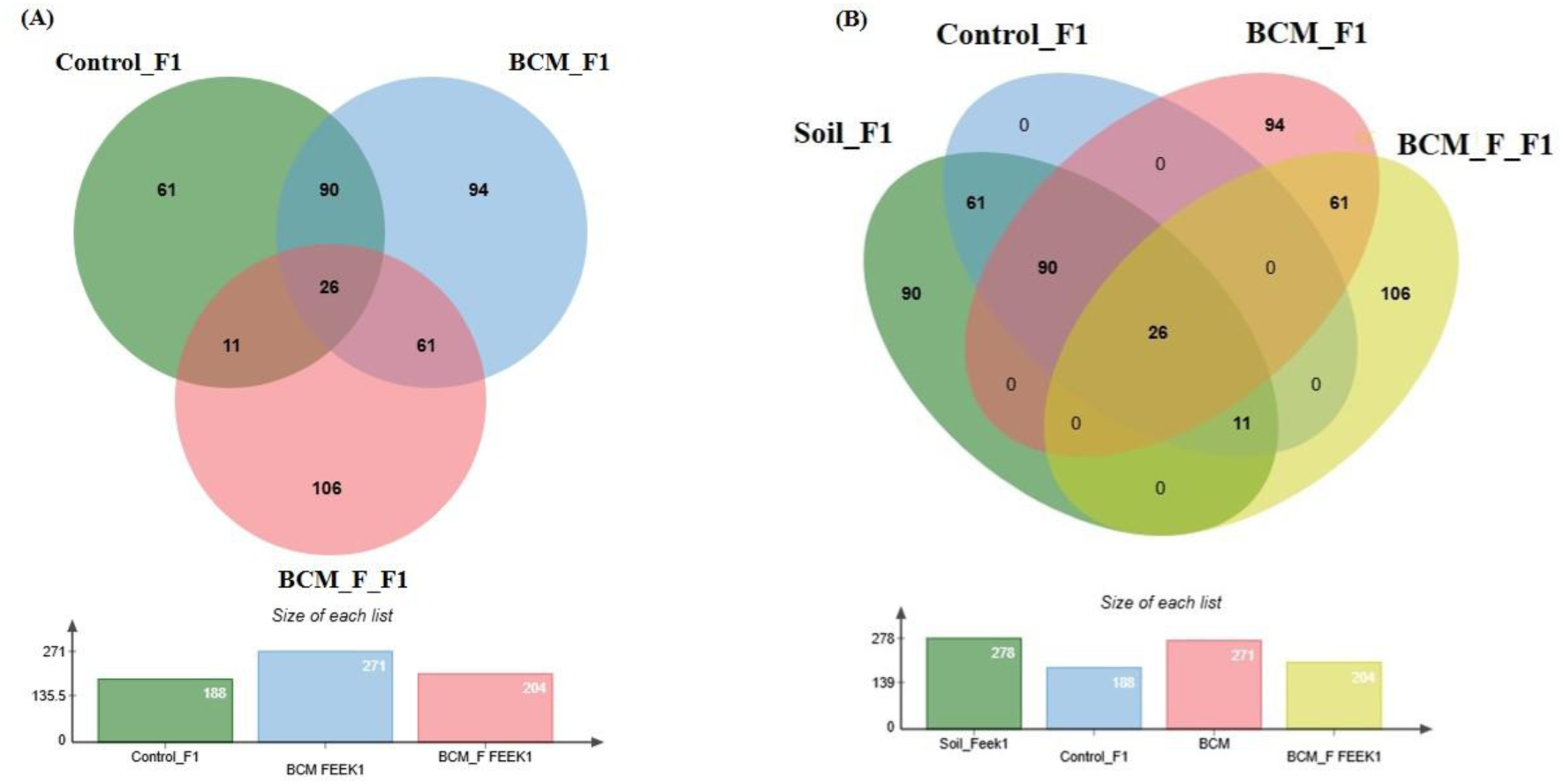
Edward’s Venn diagram illustrating the distribution of OTUs identified in wheat rhizosphere microbiota identified in the metagenomic dataset obtained after sequencing the metagenomic DNA from untreated control, BCM and BCM_F (**A**) and soil, untreated control, BCM and BCM_F (**B**). The numbers in each section represent the abundance of unique and shared microbial species at Feeks 1.0.

**Table 1:**
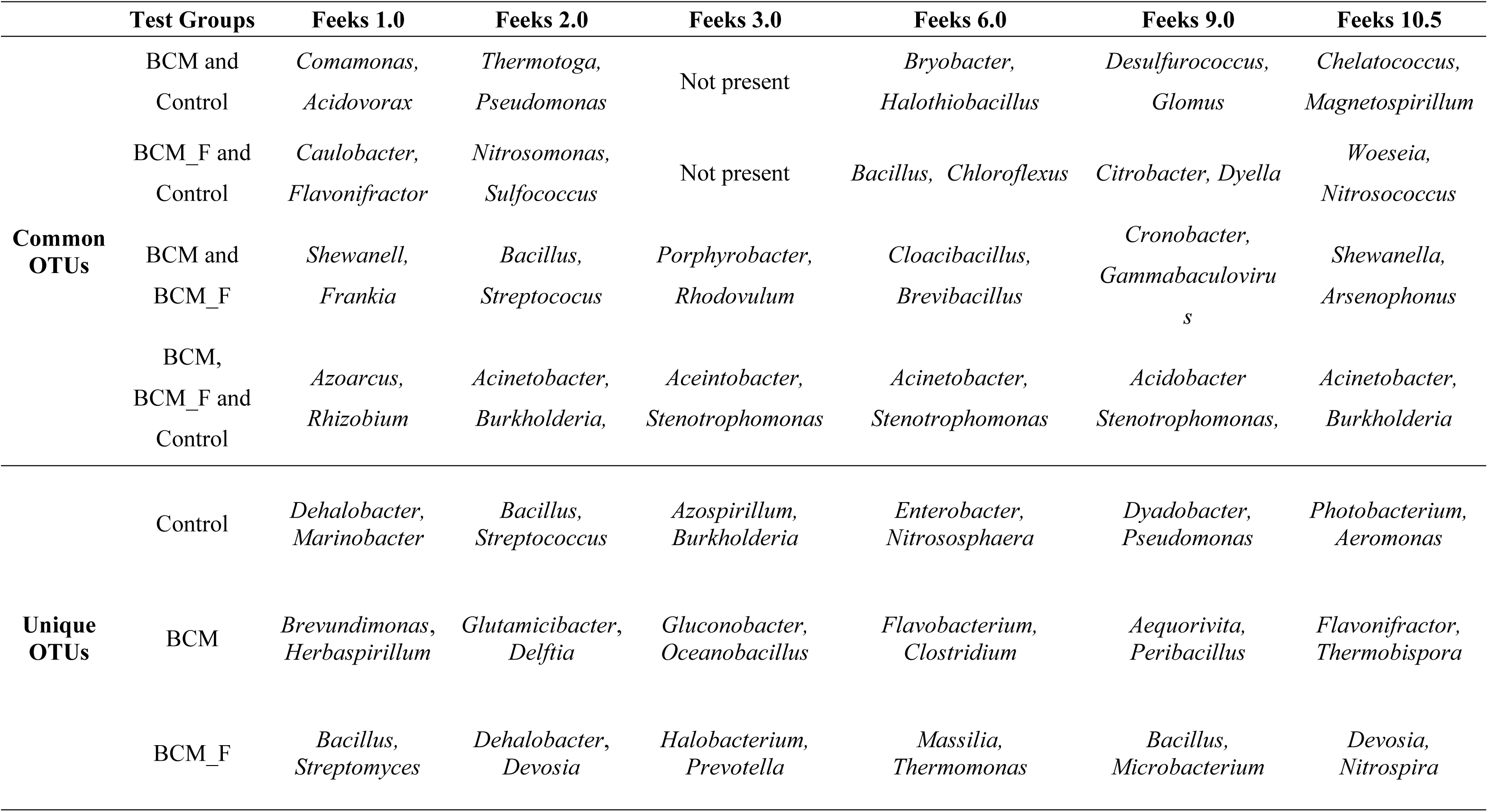
Common and unique microbial taxa associated with various test groups across wheat growth stages.

Alpha diversity analysis indicated 7.29, 6.3, and 4.97 Shannon diversity indices in BCM-treated, BCM_F-treated, and untreated wheat plants. The Shannon index reflects the abundance and evenness of species in a community, with higher values indicating greater diversity. This suggested that BCM treatment led to the highest microbial diversity, followed by BCM_F treatment, with untreated wheat plants exhibiting the lowest diversity. A comparative analysis of microbial diversity between the soil microbiota and the wheat rhizosphere microbiota at Feeks 1.0 was also conducted. The commonly shared OTUs represented 4.82% of total diversity and indicated the dominance of *Acinetobacter*, followed by *Burkholderia*. The commonly shared OTUs between control and soil represented 11.31% of total diversity and indicated the abundance of *spiribacter*, followed by *Clostridium*. The commonly shared OTUs between ‘BCM’ and ‘BCM_F’ represented 11.31% of total diversity and indicated the abundance of *Shewanella*, followed by *Ochrobactrum*. The commonly shared OTUs between ‘control’, ‘soil’, and ‘BCM_F’ represented 2.04% of total diversity and indicated the abundance of *Chryseobacterium*, followed by *Desulfovibrio.* The commonly shared OTUs between ‘control’, ‘soil’, and ‘BCM’ represented 16.69% of total diversity and indicated the abundance of *Frateuria*, followed by *Photobacterium.* The unique OTUs associated with soil represented 16.69% of total diversity and indicated the dominance of *Gemmatimonas,* followed by *Flavobacterium.* The unique OTUs associated with BCM represented 17.43% of total diversity and indicated the abundance of *Brevibacillu*s, followed by *Azorhizobium*. The unique OTUs associated with BCM_F represented 19.67% of total diversity and indicated the dominance of *Halobacterium*, followed by *Prevotella*. Commonly represented microbial OTUs in soil and wheat rhizosphere indicated soil-derived microbial acquisitions during plant growth. In contrast, the unique microbial groups indicated their origin from the seeds or external agents (anthropogenic resources) (**Figure 5B**).

### Wheat rhizosphere microbiota composition at Feeks 2.0

At Feeks 2.0 (beginning of the tillering stage), 503 OTUs were observed. The common OTUs accounted for 60.4% of the total microbial diversity. Among these, 3.379% were shared between the control and BCM, 2.78% between BCM and BCM_F, and 1.59% between BCM_F and the control. Unique OTUs associated with BCM and BCM_F represented 14.71% and 15.10% of the total observed microbial diversity (**Figure 6A**). Microbial diversity was assessed based on the presence of taxa across different Feeks stages (**Table 1**). Alpha diversity analysis indicated Shannon diversity indices of 7.85, 7.55, and 5.27 in BCM_F-treated, BCM-treated, and untreated wheat plants. These values suggested that BCM_F treatment led to the highest microbial diversity, followed by BCM_F treatment, with untreated wheat plants exhibiting the lowest diversity. Comparative microbial diversity analysis of soil microbiota and wheat rhizosphere microbiota at Feeks 2.0 was also performed. A total of 501 OTUs were identified across the samples. The OTUs commonly shared among the different groups **(Figure 6B)** accounted for 58.48% of the total microbial diversity, with *Brevibacillus* being the dominant genus, collectively representing 9.6% of the total diversity. The unique OTUs associated with BCM accounted for 14.17% of the total diversity, with *Bacillus* being the most abundant genus, followed by *Azospirillum*. Similarly, the unique OTUs associated with the untreated control samples represented 1.197% of the total diversity, with *Nitrospira* being the predominant genus, followed by *Cloacibacillus*. In the wheat rhizosphere associated with BCM_F, the unique OTUs also represented 14.17% of the total diversity, with *Steroidobacter* as the dominant genus, followed by *Bryobacter*. The unique OTUs associated with soil accounted for 2.39% of the total diversity, with *Gluconobacter* and *Halothiobacillus* being the most abundant genera.

**Figure 6:**
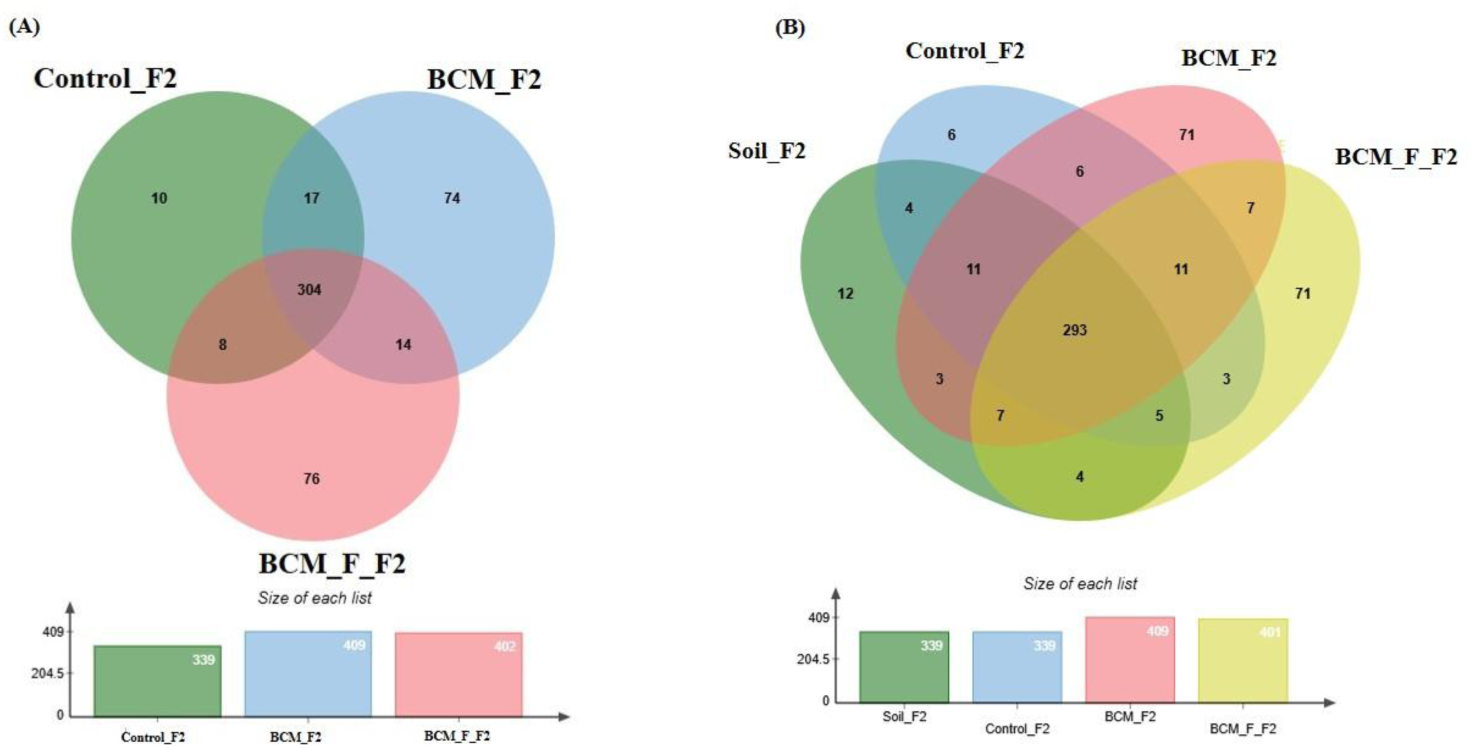
Edward’s Venn diagram illustrating the distribution of OTUs identified in wheat rhizosphere microbiota identified in the metagenomic dataset obtained after sequencing the metagenomic DNA from untreated control, BCM and BCM_F (**A**) and soil, untreated control, BCM and BCM_F (**B**). The numbers in each section represent the abundance of unique and shared microbial species at Feeks 2.0.

### Wheat rhizosphere microbiota composition and physiological role at Feeks 3.0

At Feeks 3.0 (Tillers formed), 866 OTUs were observed. The commonly shared OTUs accounted for 80.36% of the total diversity. Of the shared OTUs, 3.92% were standard between BCM and BCM_F. Unique OTUs associated with BCM, BCM_F, and the untreated control represented 7.27%, 7.27%, and 1.15% of the total diversity (**Figure 7A**). Microbial diversity was assessed based on the presence of taxa across different Feeks stages to identify unique and commonly shared microbial taxa (**Table 1**). Alpha diversity was measured in terms of Shannon diversity indices of 7.95, 7.85, and 5.94 which were observed in BCM Treated, BCM_F treated, and untreated wheat plants. Higher values indicate greater diversity; thus, BCM treatment led to the highest microbial diversity, followed by BCM_F treatment, while untreated wheat plants exhibited the lowest diversity. Comparative microbial diversity analysis of soil microbiota and wheat rhizosphere microbiota at Feeks 3.0 was also performed (**Figure 7B**). A total of 899 OTUs were observed. The common OTUs represented 52.94% of total diversity and indicated an abundance of *Massilia*, followed by *Strenotrophomonas*. Total OTUs shared by untreated control, ‘BCM’ and ‘BCM_F’ represented 24.47% of total diversity, dominated by *Pseudomonas* followed by *Acetobacter*. The commonly shared OTUs between BCM’ and ‘BCM_F represented 3.78% of total diversity and indicated an abundance of *Bacillus* followed by *Azotobacter*. The unique OTUs associated with BCM represented 1.55% of total diversity and indicated an abundance of *Flavobacterium*, followed by *Clostridium*. The unique OTUs associated with the wheat rhizosphere of BCM_F represented 42.23% of total diversity and indicated the abundance of *Sphingomonas*, followed by *Methylobacterium*. The unique OTUs associated with untreated plants represented 1.22% of total diversity and indicated the abundance of *Devosia*, followed by *Azospirillum*. The unique OTUs associated with soil represented 13.79% of total diversity and indicated the abundance of *Burkholderia*, followed by *Bradyrhizobium*.

**Figure 7:**
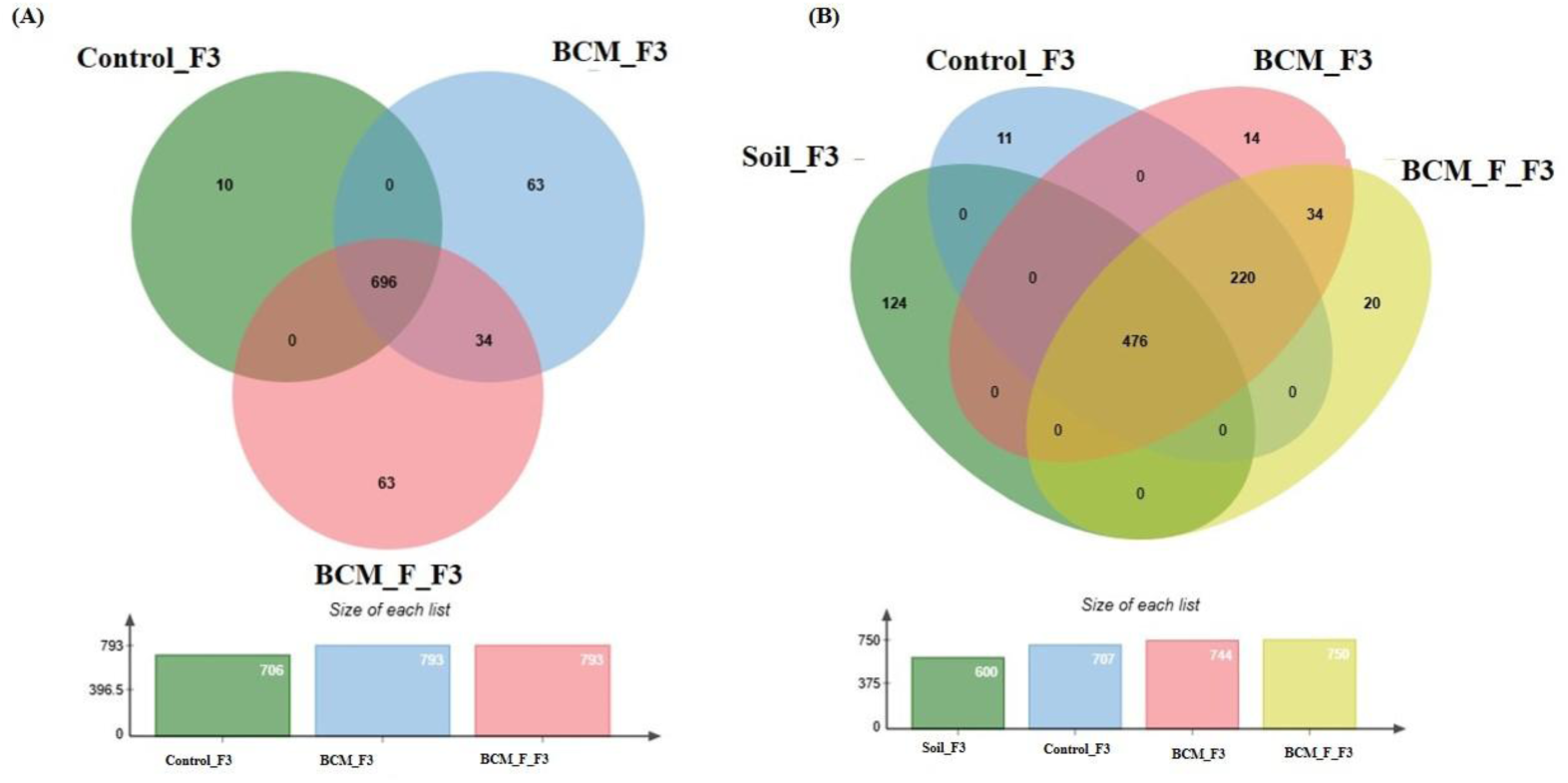
Edward’s Venn diagram illustrating the distribution of OTUs identified in wheat rhizosphere microbiota identified in the metagenomic dataset obtained after sequencing the metagenomic DNA from untreated control, BCM and BCM_F (**A**) and soil, untreated control, BCM and BCM_F (**B**). The numbers in each section represent the abundance of unique and shared microbial species at Feeks 3.0.

### Wheat rhizosphere microbiota composition at Feeks 6.0

At Feeks 6.0 (Internode formation stage), 1323 OTUs were identified. The commonly shared OTUs represented 18.82% of total diversity. The unique OTUs associated with BCM represented 32.19% of total diversity. The unique OTUs associated with BCM_F represented 30.15% of total diversity **(Figure 8A)**. Microbial diversity was evaluated by examining the presence of taxa across different Feeks stages (**Table 1**). Alpha diversity analysis indicated 8.05, 8.12, and 6.01 of Shannon diversity indices in BCM Treated, BCM_F treated, and untreated wheat plants metagenomic dataset. These values suggested that BCM treatment led to the highest microbial diversity, followed by BCM_F treatment, with untreated wheat plants exhibiting the lowest diversity.

**Figure 8:**
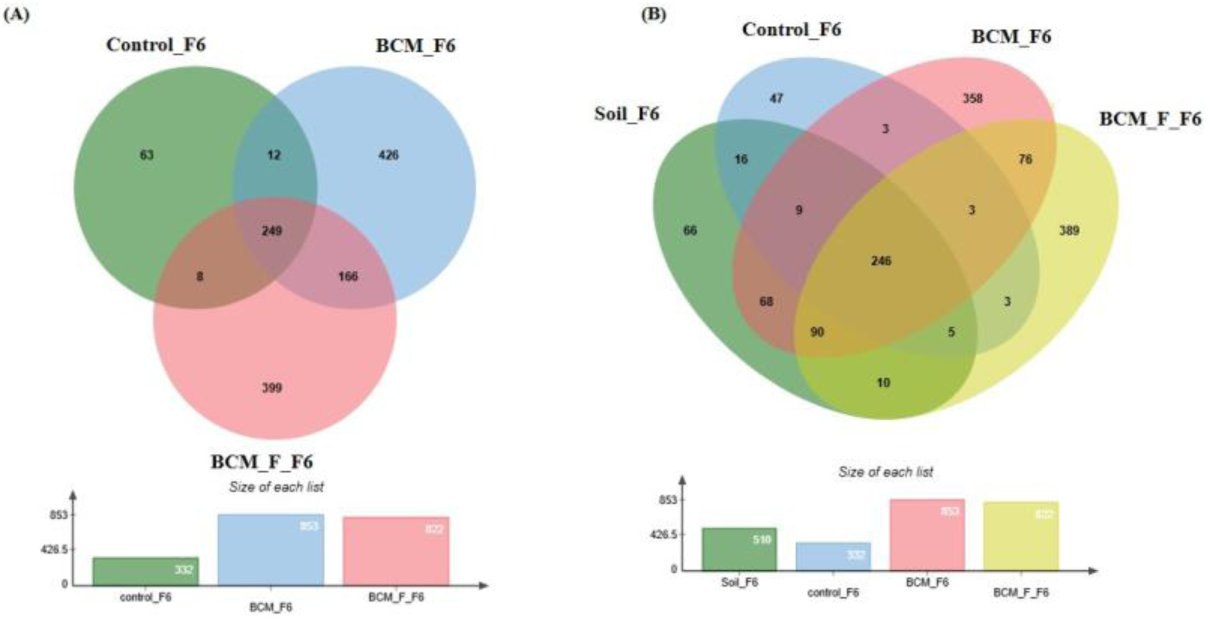
Edward’s Venn diagram illustrating the distribution of OTUs identified in wheat rhizosphere microbiota identified in the metagenomic dataset obtained after sequencing the metagenomic DNA from untreated control, BCM and BCM_F (**A**) and soil, untreated control, BCM and BCM_F (**B**). The numbers in each section represent the abundance of unique and shared microbial species at Feeks 6.0.

Comparative microbial diversity analysis of soil microbiota and wheat rhizosphere microbiota at Feeks 6.0 was performed. A total of 1389 OTUs were observed. The commonly shared OTUs represented 17.71% of total diversity **(Figure 8B)** and indicated the abundance of *Actinobacillus*, followed by *Burkholderia*. The commonly shared OTUs between ‘BCM’ and ‘BCM_F’ represented 5.47% of total diversity and indicated the abundance of *Devosia*, followed by *Bacillus*. The commonly shared OTUs between ‘control’ and ‘BCM’ represented 0.215% of total diversity and indicated the abundance of *Brachybaacterium*, followed by *Caldicellulosiruptor*. The commonly shared OTUs between ‘control’ and ‘BCM_F’ represented 0.215% of total diversity and indicated the abundance of *Streptomyces* followed by *Rhizobia*. The commonly shared OTUs between ‘soil’ and ‘BCM’ represented 4.89% of total diversity and indicated the abundance of *Chitninophaga* followed by *Glucanoacteobacter*. The commonly shared OTUs between ‘soil’ and ‘BCM_F’ represented 0.71% of total diversity and indicated the abundance of *Denitrobacterium*, followed by *Chlamydia*. The commonly shared OTUs between ‘soil’ and ‘control’ represented 1.15% of total diversity and indicated the abundance of *Desulfurococcus* followed by *Gallibacterium*. The commonly shared OTUs between ‘soil’, ‘control’ and ‘BCM_F’ represented 0.35% of total diversity and indicated dominance of *Chelatococcus* followed by *Barnesiella*. The commonly shared OTUs between ‘soil’, ‘BCM’ and ‘BCM_F’ represented 6.47% of total diversity and indicated the abundance of *Brevibacterium* followed by *Citrobacter*. The commonly shared OTUs between ‘soil’, ‘control’ and ‘BCM’ represented 0.64% of total diversity and indicated the abundance of *Gluconobacter* followed by *Aquorivita*. The commonly shared OTUs between ‘control’, ‘BCM’ and ‘BCM_F’ represented 0.215% of total diversity and indicated the abundance of *Gynuella* followed by *Desulfovibrio*. The unique OTUs associated with BCM represented 25.77% of total diversity and indicated the abundance of *Clostridium*, followed by *Streptomyces*. The unique OTUs associated with the wheat rhizosphere of untreated plants represented 3.38% of total diversity and indicated the dominance of *Acidovorax*, followed by *Spiribacte*r. The unique OTUs associated with BCM_F represented 28% of total diversity and indicated the abundance of *Flavobacterium*, followed by *Delftia*. The unique OTUs associated with soil represented 4.75% of total diversity and indicated the abundance of *Cryobacterium*, followed by *Gemmatimonas*.

### Wheat rhizosphere microbiota composition at Feeks 9.0

At Feeks 9.0 ("Ligule of flag leaf is visible" stage), 462 OTUs were observed. Commonly shared OTUs accounted for 65.8% of the total diversity, with 3.24% shared between BCM and control, 1.73% between control and BCM_F, and 2.81% between BCM and BCM_F. Unique OTUs associated with the untreated control, BCM, and BCM_F represented 2.59%, 7.53%, and 16.45% of the total diversity, respectively (**Figure 9A**). Microbial diversity was evaluated by examining the presence of taxa across distinct Feeks stages, identifying both shared and unique microbial groups (**Table 1**). Alpha diversity analysis indicated Shannon diversity indices of 5.05, 5.21, and 4.12 in BCM Treated, BCM_F treated, and untreated wheat plants. These values suggested that BCM_F treatment led to the highest microbial diversity, followed by BCM treatment, with untreated wheat plants exhibiting the lowest diversity. Comparative microbial diversity analysis of soil microbiota and wheat rhizosphere microbiota at Feeks 9.0 was performed. A total of 471 OTUs were observed. The commonly shared OTUs represented 57.74% of total diversity (**Figure 9B**) and indicated the abundance of *Acetobacter*, followed by *Azospirillum*. The commonly shared OTUs between ‘BCM’ and ‘BCM_F’ represented 5.52% of total diversity and indicated the abundance of *Gloebacter*, followed by *Bacillus*. The commonly shared OTUs between ‘control’ and ‘BCM’ represented 0.42% of total diversity and indicated the abundance of *Fermentimonas*, followed by *Barnesiella*. The commonly shared OTUs between ‘control’ and ‘BCM_F’ represented 0.63% of total diversity and indicated dominance of *Citrobacter* followed by *Edwardsiella*. The commonly shared OTUs between ‘soil’ and ‘BCM’ represented 0.84% of total diversity and indicated the abundance of *Dyadobacter* followed by *Glucanoacteobacter*. The commonly shared OTUs between ‘soil’ and ‘BCM_F’ represented 0.42% of total diversity and indicated the abundance of *Denitrobacterium* followed by *Fementimonas*. The commonly shared OTUs between ‘soil’ and ‘control’ represented 0.84% of total diversity and indicated the abundance of *Aneurinibacillus* followed by *Gallibacterium*. The commonly shared OTUs between ‘soil’, ‘control’ and ‘BCM_F’ represented 0.84% of total diversity and indicated the abundance of *Chelatophaga* followed by *Enterobacter*. The commonly shared OTUs between ‘soil’, ‘BCM’ and ‘BCM_F’ represented 1.91% of total diversity and indicated the abundance of *Brevibacterium* followed by *Massilia*. The commonly shared OTUs between ‘soil’, ‘control’ and ‘BCM’ represented 1.48% of total diversity and indicated the abundance of *Methylobacterium* followed by *Aquorivita*. The commonly shared OTUs between ‘control’, ‘BCM’ and ‘BCM_F’ represented 2.12% of total diversity and indicated the abundance of *Blautia* followed by *Nitrospora*. The unique OTUs associated with soil represented 2.33% of total diversity and indicated dominance of *Amphibacillus*, followed by *Herbaspirillum*. The unique OTUs associated with BCM represented 7.64% of total diversity and indicated dominance of *Lysobacter*, followed by *Chlorobium*. The unique OTUs associated with BCM_F represented 15.92% of total diversity and indicated dominance of *Nitrospira*, followed by *Thermotoga*. The unique OTUs associated with wheat rhizosphere of untreated plants represented 1.27% of total diversity and indicated dominance of *Thermosphaera*, followed by *Rudaea*.

**Figure 9:**
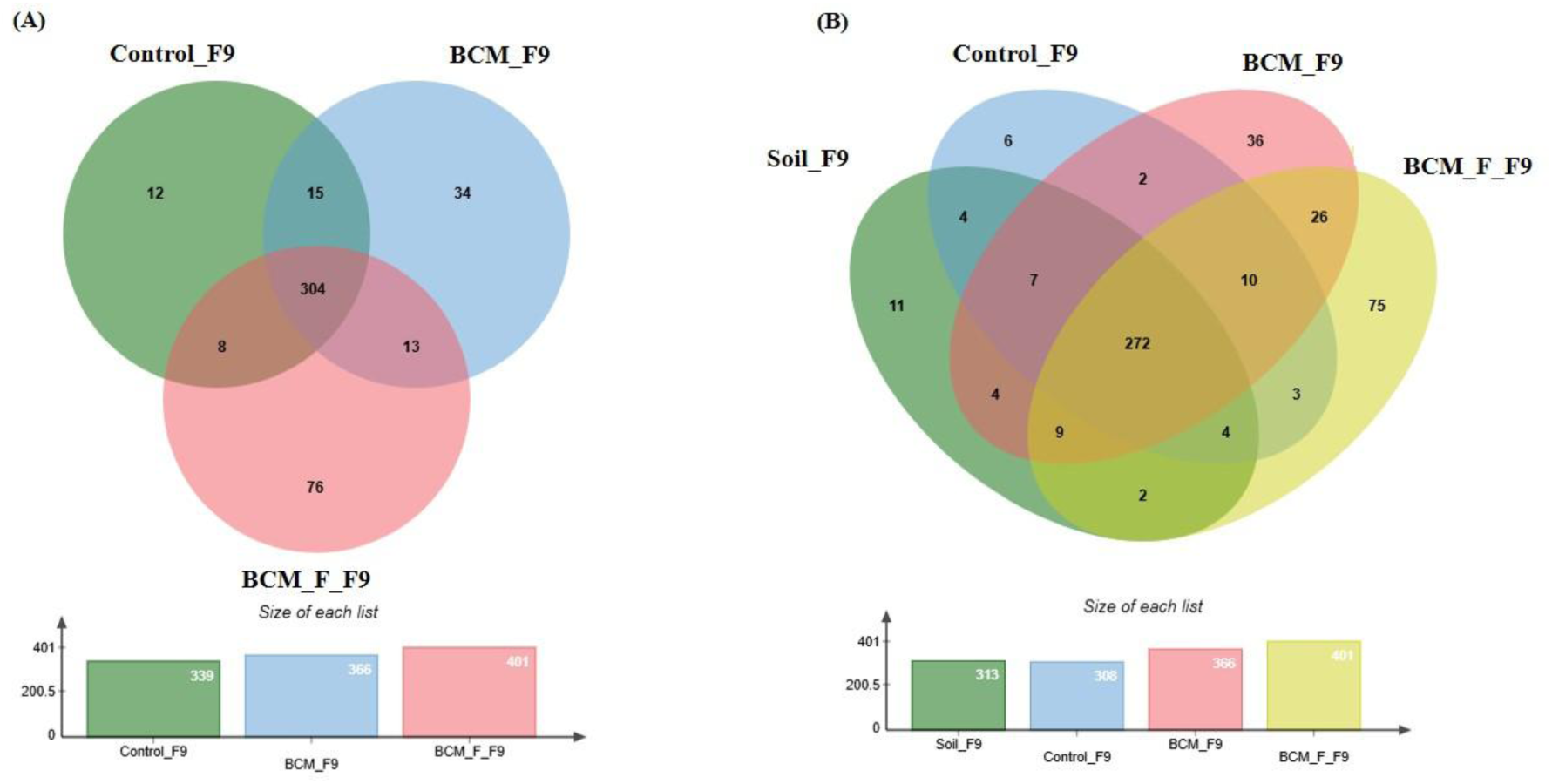
Edward’s Venn diagram illustrating the distribution of OTUs identified in wheat rhizosphere microbiota identified in the metagenomic dataset obtained after sequencing the metagenomic DNA from untreated control, BCM and BCM_F (**A**) and soil, untreated control, BCM and BCM_F (**B**). The numbers in each section represent the abundance of unique and shared microbial species at Feeks 9.0.

### Wheat rhizosphere microbiota composition and physiological role at Feeks 10.5

At Feeks 10.5 (Heading complete stage), 355 OTUs were observed. Commonly shared OTUs accounted for 82.81% of the total diversity, with 3.09% shared between BCM and control, 2.25% between control and BCM_F, and 3.09% between BCM and BCM_F. Unique OTUs associated with the untreated control, BCM, and BCM_F represented 3.09%, 1.97%, and 3.66% of the total diversity, respectively (**Table 1**, **Figure 10A**). Alpha diversity analysis indicated Shannon diversity indices of 5, 5.09, and 4 in BCM Treated, BCM_F treated, and untreated wheat plants. These suggested that BCM treatment led to the highest microbial diversity, followed by BCM_F treatment, with untreated wheat plants exhibiting the lowest diversity. Comparative microbial diversity analysis of soil and wheat rhizosphere microbiota at Feeks 10.5 was also performed. A total of 370 OTUs were observed. The commonly shared OTUs represented 76.75% of total diversity (**Figure 10B**) and indicated the abundance of *Mortierella*, followed by *Pedobacter*. The commonly shared OTUs between ‘control’ and ‘BCM_F’ represented 1.35% of total diversity and indicated the abundance of *Brucella* followed by *Streptococcus*. The commonly shared OTUs between ‘soil’ and ‘BCM’ represented 0.81% of total diversity and indicated the abundance of *Cellulomonas* followed by *Blastomonas*. The commonly shared OTUs between ‘soil’ and ‘BCM_F’ represented 1.35% of total diversity and indicated the abundance of *Granulibacter* followed by *Caulobacter*. The commonly shared OTUs between ‘soil’ and ‘control’ represented 1.08% of total diversity and indicated the abundance of *Moraxella* followed by *Dokdonella*. The commonly shared OTUs between ‘soil’, ‘control’ and ‘BCM_F’ represented 1.08% of total diversity and indicated the abundance of *Nitrosococcus* followed by *Photobacterium*. The commonly shared OTUs between ‘soil’, ‘BCM’ and ‘BCM_F’ represented 1.62% of total diversity and indicated the abundance of *Sphingomonas* followed by *Massilia*. The commonly shared OTUs between ‘soil’, ‘control’ and ‘BCM’ represented 2.43% of total diversity and indicated dominance of *Porphyrobacter* followed by *Acidiphilium*. The commonly shared OTUs between ‘control’, ‘BCM’ and ‘BCM_F’ represented 2.70% of total diversity and indicated the abundance of *Defluviimonas* followed by *Nitrospora*. The unique OTUs associated with soil represented 4.05% of total diversity and indicated the abundance of *Xanthomonas*, followed by *Thioploca*. The unique OTUs associated with untreated control represented 1.89% of total diversity and indicated the abundance of *Desulfococcus*, followed by *Frateuria*. The unique OTUs associated with BCM represented 1.08% of total diversity and indicated the abundance of *Shigella*, followed by *Hahella*. The unique OTUs associated with BCM_F represented 1.08% of total diversity and indicated the abundance of *Plantibacter*, followed by *Chromobacterium*.

**Figure 10:**
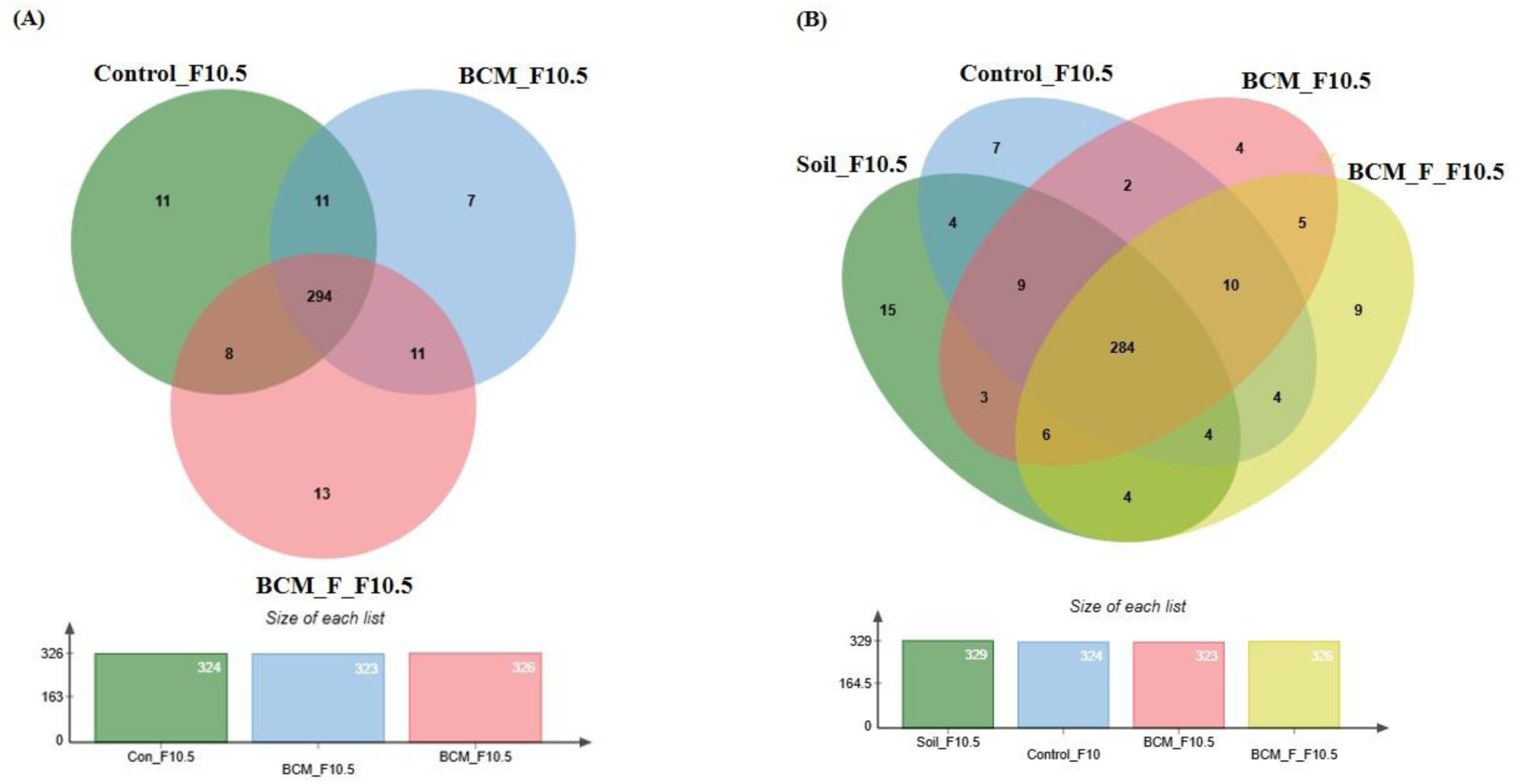
Edward’s Venn diagram illustrating the distribution of OTUs identified in wheat rhizosphere microbiota identified in the metagenomic dataset obtained after sequencing the metagenomic DNA from untreated control, BCM and BCM_F (**A**) and soil, untreated control, BCM and BCM_F (**B**). The numbers in each section represent the abundance of unique and shared microbial species at Feeks 10.5.

### Comparative analysis of Wheat rhizosphere microbiota composition at different wheat growth stages

Rhizosphere microbiota profiling showed that control soil harbors a more diverse microbial community than the wheat rhizoplane. A shift in the microbiota was observed when comparing control plants to microbial-treated plants across different growth stages. Initially, at Feeks 1.0, *Betaproteobacteria* was the most predominant class in both BCM and BCM_F. A microbial community shift was observed again at Feeks 2.0. Alphaproteobacteria **(Figure 11A)** was abundant at Feeks 2.0 followed by Betaproteobacteria **(Figure 11B)**. At Feeks 3.0, the rhizosphere microbiota of BCM and the control group were similar, with the predominance of Alphaproteobacteria **(Figure 11A)**. At the same time, a notable decrease in the abundance of Alphaproteobacteria **(Figure 11A)** was observed in BCM_F, while Betaproteobacteria was found to be abundant **(Figure 11B)**. At Feeks 6.0, BCM and BCM_F showed a shift in the microbiota from Alphaproteobacteria to Betaproteobacteria. Feeks 9.0 was predominantly dominated by a microbial community rich in Alphaproteobacteria. At Feeks 10.5, a transition from Alphaproteobacteria to Gammaproteobacteria was observed in BCM_F, while the microbiota in BCM-treated plants remained unchanged **(Figure 11)**. In the BCM-treated plants, the abundance of Gammaproteobacteria increased at Feeks 2.0 and Feek 6.0 **(Figure 11C)**. In contrast, BCM_F-treated plants exhibited a higher abundance of Gammaproteobacteria at Feeks 3.0 and Feeks 9.0 **(Figure 11C)**. Additionally, Deltaproteobacteria contributed significantly to the overall microbial diversity, with higher abundance at Feeks 3.0 in BCM-treated plants and at Feeks 6.0 in BCM_F-treated plants **(Figure 11D)**. Alongside these Proteobacteria classes, Actinobacteria was also present at all wheat Feeks stages, although in lower abundance than the Proteobacteria. Notably, the maximum abundance of Actinobacteria was observed at Feeks 2.0 and Feeks 6.0 in the BCM-treated plants, while in BCM_F-treated plants, we observed Actinobacteria abundance at Feeks 2.0 (**Figure 11E**).

**Figure 11:**
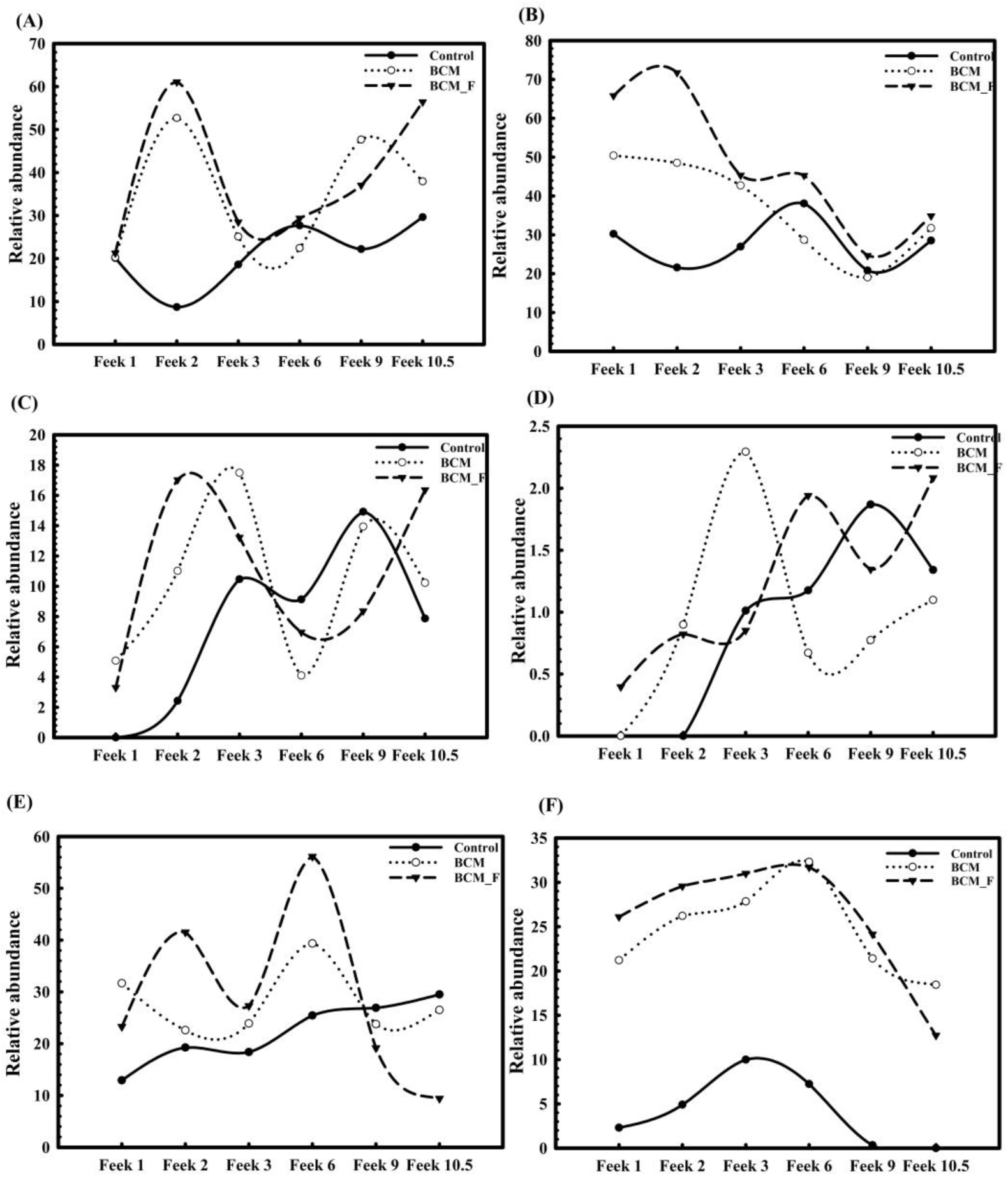
Distribution of abundant microbial classes and *Stenotrophomonas maltophilia* in the rhizosphere of plants derived from Stenotrophomonas maltophilia BCM and BCM _F treated and non-treated seeds at different Feeks (1.0, 2.0, 3.0, 6.0, 9.0, and 10.5). The figure shows the relative abundance of Alphaproteobacteria (**A**), Betaproteobacteria (**B**), Gammaproteobacteria (**C**), Deltaproteobacteria(**D**), Actinobacteria (**E**) and *Stenotrophomonas maltophilia* **(F).**

The microbial diversity across different stages of wheat growth under BCM and BCM_F treatments varied significantly, highlighting the influence of these treatments on the rhizosphere microbiota compared to untreated controls. At the emergence stage (Feeks 1.0), BCM_F treatment supported a higher proportion of unique OTUs (23.6%) dominated by *Bacillus* and *Streptomyces*, suggesting a more diverse microbial community compared to BCM (20.93%) and control (13.58%). This trend of greater microbial richness continued at subsequent stages, with BCM_F consistently promoting higher Shannon diversity indices, especially at Feeks 2.0 and Feeks 6.0. Notably, unique OTUs associated with BCM and BCM_F treatments at later stages (Feeks 6.0 and Feeks 9.0) included taxa beneficial for plant health, such as *Flavobacterium* and *Massilia* in BCM, and *Bacillus* and *Microbacterium* in BCM_F. The comparative analysis between soil and wheat rhizosphere microbiota also indicated the acquisition of soil-derived microbes throughout plant growth. At the same time, unique OTUs reflected microbial contributions from seeds or external sources. In this study, we observed significant variations in the relative abundance of *Stenotrophomonas* in treated and non-treated wheat plants. Plants treated with the microbial inoculants BCM and BCM_F exhibited significantly higher relative abundances of *Stenotrophomonas* than non-treated controls. At Feeks 1.0, *Stenotrophomonas* showed a relative abundance of 21.2% in BCM-treated plants and 26.1% in BCM_F-treated plants, while non-treated plants exhibited only 2.3% (*P < 0.001*). Similarly, at Feeks 2.0, the relative abundance of *Stenotrophomonas* was significantly higher in treated plants (26.2% in BCM and 29.56% in BCM_F) compared to the non-treated plants (4.9%) (*P < 0.001*). Interestingly, *Stenotrophomonas* was also more abundant in the soil (22.36%) than in the control group, indicating its strong presence in the environments **(Figure 11F)**. As the plants progressed through different growth stages, the relative abundance of *Stenotrophomonas* continued to increase in both BCM and BCM_F- treated groups. At Feeks 3.0, the abundance of *Stenotrophomonas* was notably higher in the treated plants (27.8% in BCM and 31% in BCM_F) than in the control (16.85%) (*P < 0.001*). Moreover, the class Gammaproteobacteria showed a higher abundance of *Stenotrophomonas* in treated plants (32.3% in BCM and 31.69% in BCM_F) compared to the non-treated plants (19.98%) at Feeks 6.0 **(Figure 11F)**. However, at later stages (Feeks 9.0), a slight decrease in *Stenotrophomonas* abundance was observed in both BCM (21.4%) and BCM_F (24.15%) treatments, as compared to the previous stages, though the abundance remained higher than in the untreated control (7.5%) (P < 0.001). Additionally, *Stenotrophomonas* 16S rRNA gene sequences were consistently present throughout all six growth stages in BCM and BCM_F- treated wheat plants, with notably higher abundance than untreated plants **(Figure 11F)**. These results indicate the stability of *Stenotrophomonas maltophilia* BCM and BCM_F across plant growth stages, which is essential to avail their plant growth promotion properties for sustainable growth enhancement.

### Sugar contents, phosphatase activity, and nitrogen assimilation properties in wheat rhizosphere at various Feeks

The rhizosphere of wheat plants from seeds pre-treated with *Stenotrophomonas maltophilia* BCM and BCM_F consistently showed significant improvements in various physiological parameters compared to the untreated seeds. During the growth stages (Feeks 1, 2, 3, 6, 9, and 10.5), the total sugar (**Supplementary Figure SF 2A)** and reducing sugar content (**Supplementary Figure SF 2B)** in the rhizosphere was notably higher in the pre-treated plants, with increases ranging from approximately ∼2.1 to ∼4.38 times for total sugars and ∼0.10 to ∼1.51 times for reducing sugars (**Supplementary Figure SF2)**. The pre-treatment with *Stenotrophomonas maltophilia* BCM and BCM_F enhanced nitrogen assimilation in the rhizosphere, as indicated by the increased nitrate reductase activity (**Supplementary Figure SF 3A)**. This activity was boosted by as much as ∼8.25 times at Feeks 6 and showed consistent improvements at other stages, ranging from ∼1.33 to ∼8.25 times higher than in untreated seeds. Similarly, extracellular alkaline phosphatase activity was significantly elevated in the pre-treated plants, with increases varying from ∼3 to ∼7.93 times across different stages (**Supplementary Figure SF 3B)**. These findings suggest that the pre-treatment improved nutrient cycling, particularly in terms of sugars and phosphorus, and also contributed to enhanced nitrogen uptake, ultimately supporting better growth and development of the wheat plants (**Supplementary Figure SF2 and SF3**).

### Assessment of alteration in phenotypic variation of wheat plants

Wheat plants treated with *Stenotrophomonas maltophilia* BCM and *Stenotrophomonas maltophilia* BCM_F showed a significant increase in the number of tillers (*P=0.0024*), number of leaves per plant (*P=0.0003*), spike length (*P=0.00012*), number of spikes per plant (*P=0.00036*), number of spikelets per plant (*P=0.0001*), grain weight per 1000 grains (*P=0.0031*) and grain yield (*P=0.0002*) (**Table 2**). These findings suggested that *Stenotrophomonas maltophilia* BCM and BCM_F may enhance wheat productivity by promoting both vegetative growth (e.g., more tillers and leaves) and reproductive success (e.g., longer spikes, more spikelets, and higher grain weight), leading to higher overall grain yields. This could have important implications for sustainable agriculture, particularly as these microbial isolates may reduce the need for chemical fertilizers and enhance crop performance under varying environmental conditions.

**Table 2:** Assessment of plant productivity features in *Stenotrophomonas maltophilia* treated seeds in comparison to untreated. All the experiments were performed in triplicates. Values in the table represent the mean and S.D.

| Productivity phenotype | WC-306 Plants | WC-306 plants treated with *Stenotrophomonas maltophilia* BCM | WC-306 plants treated with *Stenotrophomonas maltophilia* BCM_F |
| --- | --- | --- | --- |
| Leaves per plant | 12.34±0.57 | 44.52±1 | 33.57±.257 |
| Number of tillers | 3.34±0.57 | 5.67±0.57 | 4±.31 |
| Number of spike/plant | 26.3±1.309 | 50.67±1.52 | 42.44±1.03 |
| Spike length (cm) | 15.46±0.41 | 23.5±0.96 | 21.2±0.6 |
| Number of spikelets per plant | 30.34±0.57 | 55.6±1.52 | 45.67±0.152 |
| Grain weight (g) | 30.23±0.37 | 56±0.201 | 51.5±0.08 |
| Grain yield (Kg/acre) | 3267 | 64255.6 | 56005 |

## Discussion

Wheat, a staple crop critical to global food security (Tilman et al., 2011), faces increasing threats from biotic and abiotic stressors that jeopardize production. These include rising soil salinity, declining soil fertility (Gamage et al., 2023), the emergence of phytopathogens (Newman and Derbyshire, 2020), and the adverse effects of climate change (Srinivas et al., 2019). To address these challenges, there is an urgent need to identify microbial agents that can potentially enhance wheat’s resilience to biotic stresses, such as pathogens, and abiotic stresses, such as salinity, thereby improving crop productivity (Sharma et al., 2021; Mehmet Tuğrul, 2020). The rhizosphere is a rich source of biofertilizers due to its dense population of beneficial microorganisms that interact with plant roots (Sharma et al., 2021). These inhabiting microbes include nitrogen-fixing bacteria, mycorrhizal fungi, and phosphate-solubilizing bacteria, enhancing nutrient availability, improving soil structure, and promoting plant growth (Koza et al., 2022). The root exudates plants release serve as a continuous energy source for these microorganisms, enabling their prolonged establishment (Sharma et al., 2021). As a result, rhizosphere-derived biofertilizers can improve nutrient uptake, support plant health, and reduce reliance on chemical fertilizers (Sharma et al., 2024b). This study aimed to explore the wheat rhizosphere microbiota in quest of microbial candidates that can promote plant growth, inhibit pathogen infections, enhance nutrient assimilation, and improve resistance to environmental stressors. Exploration of wheat rhizosphere identified two potent microbial strains with biocontrol and biofertilizing properties. The 16S rRNA gene of BCM and BCM_F shared similarity with *Stenotrophomonas maltophilia*, and BCM was already characterized for its fertilization and bio-control potential in our previous study (Sharma et al., 2024a). Notably, the BCM_F strain also demonstrated antifungal activity against *Fusarium oxysporum*, with its inhibitory effect being more potent than that of the BCM strain. BCM_F is an effective biocontrol agent and biofertilizer due to its remarkable growth-promoting abilities and resilience under challenging environmental conditions. BCM_F can stimulate plant growth by enhancing nutrient availability via phosphate and ammonium solubilization, promoting root development, and producing growth-promoting substances such as IAA. Additionally, BCM_F exhibits exceptional tolerance to high salinity, metal toxicity, drought, and oxidative stress, common stressors in many agricultural environments (Sharma et al., 2024a). This ability to thrive under such conditions allows BCM_F to promote plant growth and protect plants from stress-induced damage. As a result, BCM_F serves dual functions, such as improving soil fertility and plant health as a biofertilizer while also acting as a biocontrol agent by suppressing soil-borne pathogens such as *Fusarium oxysporum* and enhancing plant resistance to diseases, making it a valuable tool for sustainable agriculture. Results of the field study showcased the significant improvement in plant growth parameters and crop yield, indicating their efficacy as biofertilizers and biocontrol agents beyond experimental conditions. Furthermore, rhizosphere metagenomics profiling identified a higher microbial community abundance in the wheat rhizosphere in treated plants compared to non-treated plants. Additionally, a shift in the microbiota was observed between control and microbial-treated plants at different growth stages. Microbial community dynamics and higher diversity could result from plant growth, increased root density (Ma et al., 2021), or microbial preincubation (Sharma et al.,2024b). The increased microbial diversity also enriched nitrogen-fixing strains such as *Azotobacter*, *Rhizobium*, *Bradyrhizobium*, and *Pseudomonas*, which are well known for enhancing nitrogen availability through processes like nitrogen fixation (Saeed et al., 2021). Additionally, the observed increase in phosphate solubilization at these Feeks may be due to the proliferation of phosphate-solubilizing microbes, including *Enterobacter*, *Penicillium*, *Streptomyces*, and *Bacillus*, which are capable of converting insoluble phosphorus into soluble forms accessible to plants (Saeed et al., 2021). *Stenotrophomonas maltophilia* BCM and BCM_F specific 16S rRNA gene sequences were observed in all six growth stages of the wheat rhizosphere in microbial-treated plants and untreated plants with lower abundance. The tillering stage in wheat development has agronomic importance (Liu et al., 2023). However, this stage is also very prone to pathogen attacks. During wheat’s tillering and internode formation stages, the impact of *Rhizoctonia solani* and *Fusarium oxysporum* can be quite detrimental to plant growth and development (Mulk et al., 2022; Castro et al., 2019). In the tillering stage, *Rhizoctonia solani*, a soil-borne pathogen, causes root rot and damping-off, impairing the establishment of new tillers by damaging the root system (Sturrock et al., 2015). This would result in reduced tiller number, poor root development, and weakened plant vigor. Both pathogens hinder the plant’s ability to thrive during this critical growth phase, leading to lower yield potential. Interestingly, we observed an increased abundance of *Stenotrophomonas maltophilia* in Feeks 1.0, Feeks 3.0, and Feeks 6.0. *Stenotrophomonas maltophilia* has been characterized by biocontrol properties against *Rhizoctonia solani* and *Fusarium oxysporum* (Sharma et al., 2024a). The presence of *Stenotrophomonas maltophilia* BCM and BCM_F in wheat rhizosphere will ensure protection against fungal infection. Microbes extending protection against these phytopathogens are known to improve crop yield (Mulk et al., 2022). So, enhanced wheat crop yield observed with microbial pre-inoculation could be attributed to *Stenotrophomonas maltophilia* presence across plant growth stages.

Microbial preincubation is known to modulate rhizosphere microbiota composition (Sharam, et al., 2024b). Likewise, a shift in microbiota composition was observed in the rhizosphere of plants originating from *Stenotrophomonas maltophilia* inoculated seeds. *Dyodobacter*, *Clostridium*, *Sphingobacterium*, *Vicinamicibacter*, *Sphingomonas* were exclusively associated with Feeks 3.0. *Dyadobacter* sp. was found to be responsible for nitrogen fixation in nitrogen- deficient conditions and promoting plant growth (Kumar et al., 2018). *Clostridium sp.* exhibits activity across diverse environmental conditions, contributing to agroecological benefits by promoting plant growth (Doni et al., 2014). *Clostridium* sp. can facilitate biological nitrogen fixation and phosphate solubilization processes, thereby enhancing the bioavailability of these essential nutrients to plants (Timofeeva et al., 2023). Some of the *Sphingobacterium* sp. such as *Sphingobacterium anhuense* and *Sphingobacterium pakistanensis. Sphingobacterium* is a soil bacterium that has shown potential in promoting plant growth. It was identified to solubilize essential nutrients such as phosphorus and produce phytohormones such as gibberellins and auxin. *Vicinamicibacter* can solubilize phosphate, transforming insoluble phosphorus compounds into soluble forms that plants can absorb, thus improving phosphorus availability, a crucial nutrient for root development and energy metabolism (Pan and Cai, 2023). Beyond nutrient enhancement, *Vicinamicibacter* may contribute to plant health by inducing systemic resistance to pathogens, bolstering the plant’s defense mechanisms, and improving overall stress tolerance (Heil and Bostock, 2002). Similarly, *Sphingomonas* strains contribute to plant growth promotion while also bolstering plant resistance to pathogens (Mazoyon et al., 2023). *Sphingomonas trueperi*, *Sphingomonas sediminicola* Dae20 have been shown to enhance plant defense mechanisms, improving resistance to a broad spectrum of pathogens, including fungi and bacteria (Mazoyon et al.,2023). At Feeks, 6.0, 16S rRNA genes belonging to *Flavobacterium* were exclusively observed in microbial-treated groups at this Feeks. *Flavobacterium* species have been shown to positively impact plant health and development by promoting growth, controlling diseases, and enhancing tolerance to abiotic stress (Seo et al., 2024). Feeks 9.0 and 10.5 also had more microbial diversity than the control plants. The study’s findings demonstrate that the observed shift in microbial communities at Feeks 3.0 and 6.0 enhances plant growth performance under abiotic stressors, such as drought and salt stress, while simultaneously providing resistance to pathogenic organisms. In parallel, we recorded improvements in key plant health indicators, including total sugar, reducing sugar, and nitrite concentrations at Feeks growth stages 3.0 and 6.0. From this, the enhanced microbial diversity at Feeks 3.0 and 6.0 contributed significantly to improved nutrient cycling, with beneficial impacts on nitrogen and phosphorus availability, ultimately supporting plant growth and soil fertility.

Rhizosphere microbiota comparison with soil microbiota unveiled the source of the rhizospheric microbiota’s origin. The majority of microbial members in rhizosphere microbiota are the same as observed in soil, indicating their origin. However, several unique OTUs were observed for both soil and rhizosphere samples. Unique OTUs in soil indicated that all microbial members cannot colonize. Host plant genotype is a major regulating factor in defining microbiota composition (Wu et al., 2025). This differential selection pressure could be a reason for it. However, unique OTUs in the rhizosphere of *Stenotrophomonas maltophilia-*treated groups generate curiosity about their origin. *Pseudomonas*, *Bacillus*, and *Staphylococcus* are identified from air samples (Chen et al., 2024), so air might have contributed the same. Additionally, *Brachybacterium (*Mounier et al., 2017), *Caldicellulosiruptor* (Byrne et al., 2021), *Glucanoacetobacter* (Pallucchini et al., 2024)*, Chelatococcus* (Yoon et al., 2008)*, Gemmatimonas* (Bandopadhyay and Shade, 2024) were identified in water ecosystem, so these microbial groups might have attained in rhizosphere during irrigation. *Barnesiella* is usually present in the human colon (Daillère et al., 2016). It can be concluded that wind, wind-blown soil, and water through irrigation could be the possible sources of these unique OTUs. Additionally, the present study defines the stability of *Stenotrophomonas maltophilia* throughout plant growth stages and explores their influence on rhizosphere microbiota. Collectively, our data underscore the potential of *Stenotrophomonas maltophilia* BCM and BCM_F as efficacious biofertilizers beyond their biocontrol activity against *Rhizoctonia solani* and *Fusarium oxysporum*. his strategy presents a cost-effective and ecologically sustainable alternative to conventional chemical fertilizers, facilitating improved nutrient bioavailability to plants and promoting plant growth under stressful conditions.

## Author Contributions

NSC designed the study and experiments. NSC, PS, and RP wrote the manuscript. PS carried out the experiments. NSC, PS, and RP analyzed the data. All authors edited the manuscript and approved the final draft of the manuscript.

## Acknowledgment

The authors acknowledge CSIR-Institute of Genomics and Integrative Biology, New Delhi, India for DNA sequencing facility.

## Data Availability Statement

The 16S rRNA gene sequence datasets generated in this study were deposited at NCBI with SRA accession ID PRJNA1136665 (https://www.ncbi.nlm.nih.gov/sra/PRJNA1136665).

