## Supplementary Figures for "*Stenotrophomonas maltophilia* promotes wheat growth by enhancing nutrient assimilation and rhizosphere microbiota modulation"

**Running Title:** *Stenotrophomonas maltophilia* for sustainable agriculture

**Number of Words:** 10859
**Number of Figures:** 11
**Number of Tables:**02

**Supplementary Figure SF1:** Microbial abundance at phylum level of taxonomy observed in the rhizosphere of microbial treated and non-treated plants at different Feeks (1.0, 2.0, 3.0, 6.0, 9.0, 10.5). Here 1-316 represents different phylums represented as 1: Proteobacteria, 2: Firmicutes, 3: Actinobacteria, 4: Cyanobacteria, 5: Bacteroidetes, 6: Acidobacteria, 7: Chloroflexi, 8: Synergistetes, 9: Verrucomicrobia, 10: Tenericutes, 11: Planctomycetes, 12: Gemmatimonadetes, 13: Nitrospirae, 14: Deinococcus-Thermus, 15: Thermotogae, 16: Chlorobi

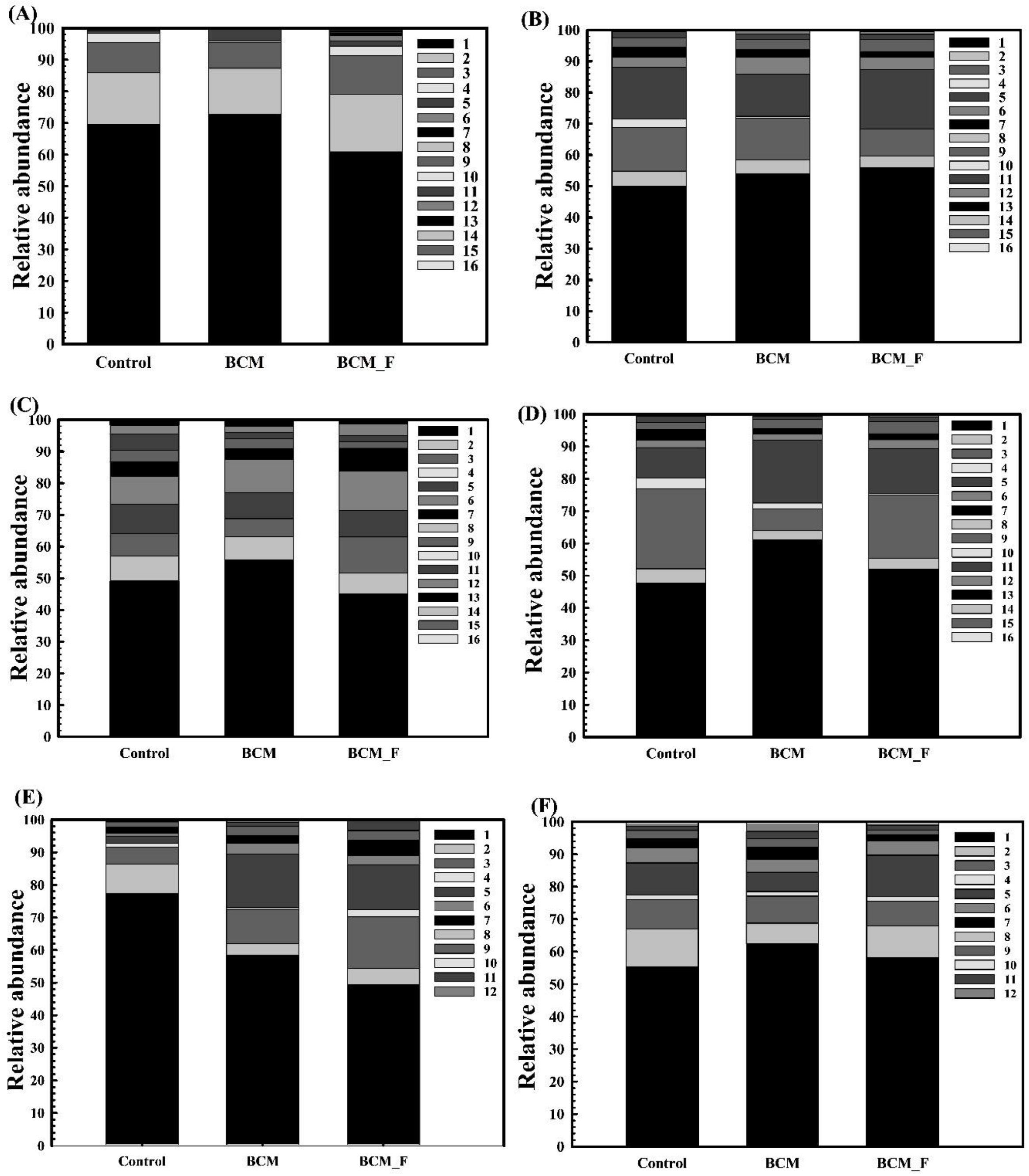

**Supplementary Figure SF2:** Total sugar content observed at different Feeks (1.0, 2.0, 3.0, 6.0, 9.0, 10.5) in the presence and the absence of microbial inoculants BCM and BCM_F (**A**). Reducing sugar was estimated using DNS assay at different Feeks (1.0, 2.0, 3.0, 6.0, 9.0, 10.5) was assessed in the presence and the absence of microbial inoculants BCM and BCM_F (**B**). The experiment was carried out in triplicates. Plotted values are the mean of triplicate readings and their observed standard deviation.

(A)

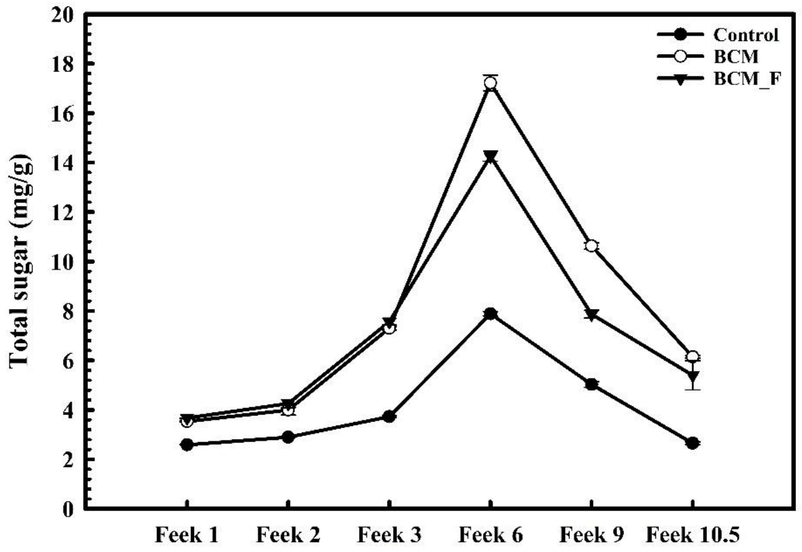


(B)

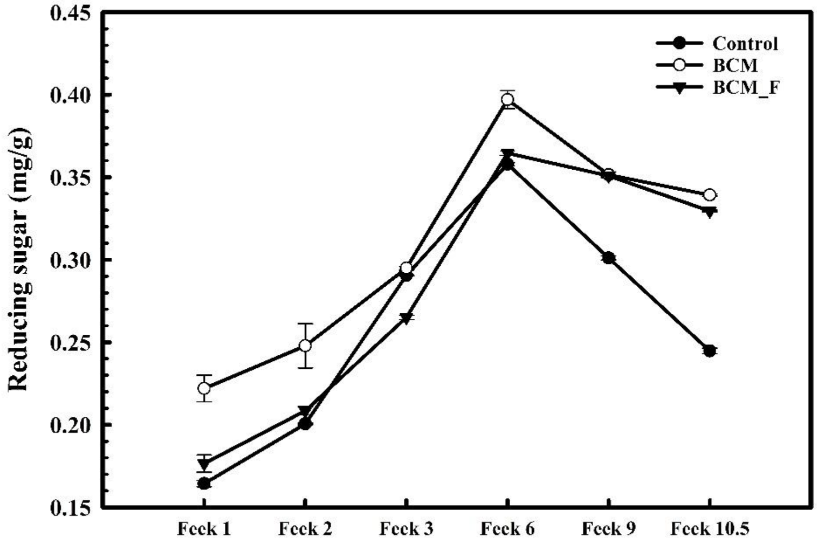

**Supplementary Figure SF3:** Assessment of nitrate reductase activity at different Feeks (1.0, 2.0, 3.0, 6.0, 9.0, and 10.5) in the presence or absence of microbial groups *Stenotrophomonas maltophilia* BCM and BCM _F (**A**). Assessment of alkaline phosphatase activity at different Feeks (1.0, 2.0, 3.0, 6.0, 9.0, 10.5) in the presence or absence of BCM and BCM_F(**B**). Experiments were carried out in triplicates Plotted values are the mean of triplicate readings and their observed standard deviation.

(A)

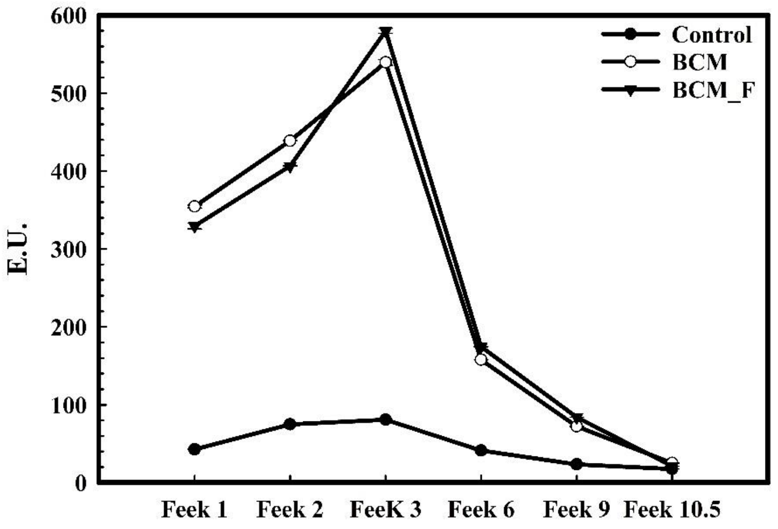


(B)

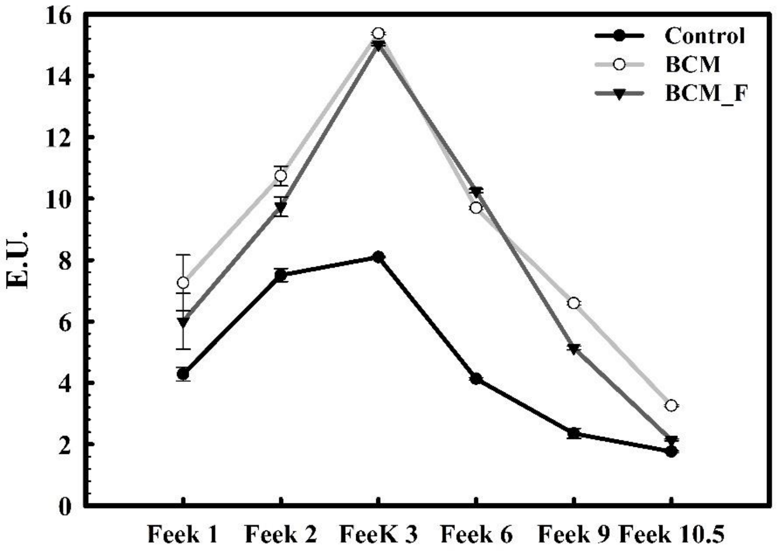
